# SynAPSeg: A novel dataset and image analysis framework for deep learning-based synapse detection and quantification

**DOI:** 10.64898/2026.03.12.711395

**Authors:** Pascal Schamber, Sahana Darbhamulla, Molly Boyer, Madison Pelletier, Helene Hartman, Olivia Friedman, Shiyu Zhang, Allison Blais, Seyun Oh, Haining Zhong, Alexei M Bygrave

## Abstract

Synapses are the fundamental units of neural computation, yet quantifying their organization across circuit-level scales remains a critical bottleneck in neuroscience. While advances in fluorescent labeling and imaging can generate vast datasets, analysis is often the limiting factor. Several deep learning-based tools have been proposed to ameliorate these issues. However, existing applications primarily focus on dendritic spines and lack robust solutions for segmenting synaptic puncta in dense tissue preparations. To address this, we introduce SynAPSeg, which encompasses an open-source framework for deep learning-based analysis and, to the best of our knowledge, the first large-scale, publicly available instance segmentation dataset specifically curated for synaptic puncta. We use this dataset to train deep learning models that reach the performance of human experts across a unique benchmark dataset. SynAPSeg integrates these models into an interactive interface, with support for multi-dimensional data, enabling fully automated segmentation and quantification pipelines alongside an annotation module for refinement and validation. We demonstrate the framework’s scalability by performing the first comprehensive mapping of nearly 4 million excitatory postsynaptic PSD95 puncta within inhibitory interneurons across the dorsal hippocampus, revealing regional differences in synapse properties. Finally, we show SynAPSeg’s utility for 3D quantification by applying these models to study aging-associated synaptic changes in CA1 parvalbumin (PV)-positive inhibitory neurons. Through this approach, we uncover a reduction in PSD95 density along PV dendrites in the aged CA1, indicating reduced glutamatergic recruitment of PV neurons which could contribute to age-related cognitive decline. Collectively, these results demonstrate that SynAPSeg provides a scalable solution for comprehensively studying synaptic architecture in health and disease.

## Introduction

Synapses are fundamental units of computation in the brain; their strength and connectivity define the functional architecture of neural circuits and underlying plasticity. Quantifying synaptic changes at scale is essential for bridging molecular and circuit-level neuroscience yet it remains a significant technical challenge. While electron microscopy remains the gold standard for studying ultrastructure, laser scanning microscopy (LSM) offers scalability, and the capacity for multiplexed imaging of synaptic proteins across broad neural circuits. LSM also benefits from recent innovations in genetic mouse models and the development of intrabodies, which enable the fluorescent labeling of endogenous synaptic proteins with enhanced signal-to-noise ratios (SNR) and cell-type specificity compared to traditional immunohistochemistry (IHC) ^1–4^. Collectively, this progress shifts the bottleneck in achieving scale from image acquisition to quantification.

While manual annotation is widely accepted as the gold standard for quantifying synapses, its prohibitive time requirements make it impractical for large datasets. In addition, recent studies have shown high inter-rater and intra-rater variability even among experts, challenging the idea that manual annotations reliably reflect an absolute ground truth ^4–6^. This ambiguity can be partially attributed to the small size of synapses; as nanoscale structures they often approach the resolution limits of LSM making precise delineation of object boundaries challenging ^7,8^. Consequently, annotations from multiple experts are frequently pooled to generate a consensus, though further exacerbate resource demands ^6,8–10^. Automated thresholding-based approaches have been employed to mitigate subjectivity. However, despite the advances made by these methods, they often provide fragile solutions that require parameters to be manually tuned per-image to accommodate varying SNR and illumination levels ^11–14^, effectively reintroducing the bottleneck and reproducibility issues they were meant to combat. Conventual techniques also typically struggle to resolve object boundaries when signal is dense, especially in volumetric (3D) contexts, as they depend on heuristic-based methods like flood-filling or watershed segmentation to delineate individual objects that are prone to merging errors, leading to under-segmentation, and compromise the ability to analyze the features of individual synaptic structures ^7,11,14,15^.

Deep learning-based methods have been applied to address similar fluorescent imaging challenges in other areas of biology ^16–20^, and recently, to dendritic spine segmentation. DeepD3 for example, can accurately segment fluorescent cell fill signal to detect dendritic spines in challenging in vivo domains ^6^. Others have shown incorporating image restoration techniques can further improve performance ^8,21^. While these approaches work well for sparse dendritic spines, the underlying U-Net-like architecture performs semantic segmentation and does not resolve individual objects. Instead, they must rely on conventional heuristic-based methods to separate touching objects and thus face similar challenges on dense labels. In addition, the punctate signals produced by fluorescently labeled synaptic proteins are morphologically distinct from cell filled spines. This unique detection challenge potentially limits the analysis of various synaptic puncta including somatic synapses, most inhibitory synapses, as well as excitatory inputs onto inhibitory neurons which typically form shaft synapses along dendrites rather than dendritic spines ^22^.

The largest challenge in developing robust deep learning-based approaches for specialized domains remains a lack of publicly available training data ^23,24^. It has been repeatedly shown that deep learning models struggle with data far outside the training distribution ^6,8^, therefore without dedicated training datasets synaptic puncta segmentation will remain limited. To the best of our knowledge, the DeepD3 dataset is the only significant collection of manually annotated training data in this domain. While we generally lack specialized end-to-end deep learning tools capable of segmenting individual puncta, generalist architectures such as StarDist ^25–27^, Cellpose ^28–30^, and MASK R-CNN ^31^ have been widely applied for instance segmentation across biological domains as well as in dendritic spine segmentation ^32^. However, the coding expertise required to utilize and properly implement these tools can create a technical barrier, especially for 3D data, as popular image analysis tools such as ImageJ lack full support even for popular techniques like StarDist.

To facilitate deep-learning-based synaptic quantification, we introduce SynAPSeg, encompassing both open-source datasets and a framework for image analysis. The datasets span a diverse range of labeling modalities to promote generalizability and represent, to our knowledge, the first large-scale, publicly available instance segmentation dataset for synaptic puncta. We use this dataset to evaluate the performance of several recent algorithms for synapse segmentation and train custom deep learning-based models – finding that custom-trained StarDist architectures show the highest performance on 2D and 3D instance segmentation tasks. We benchmark these models using a novel multi-rater dataset and demonstrate that they reach the level of human experts. These models are made accessible through the SynAPSeg framework, providing a user-friendly interface that combines segmentation, validation, and quantification into a single workflow. We demonstrate the scalability of this platform by applying SynAPSeg to generate the first comprehensive mapping of excitatory postsynaptic densities—through imaging of PSD95—on inhibitory interneurons across the dorsal hippocampus. Lastly, we demonstrate the utility of SynAPSeg for robust 3D quantification, replicating findings that PV intensity correlates with the number of excitatory synapses, as well as uncovering an age-related reduction in PSD95 puncta density along PV dendrites.

## Results

### Dataset creation and method valuation

In contrast to the extensive development of deep learning architectures for synapse detection^6,8,10,23,33,34^, the field generally lacks publicly available data to train such models. To address this gap, we generated diverse training datasets containing pixel-precise manual annotations for over 4300 2D and 1200 3D putative synaptic puncta (**Fig 1A**). To provide broad utility, the datasets encompass different resolutions, sample preparations, synaptic markers, labeling methods, SNR, and morphologies (detailed in **Supplementary Table 1**). Given significant progress in spine detection, one focus of this dataset was improving the segmentation of *aspiny* postsynaptic densities, commonly referred to as “shaft synapses”.

**Figure 1.**
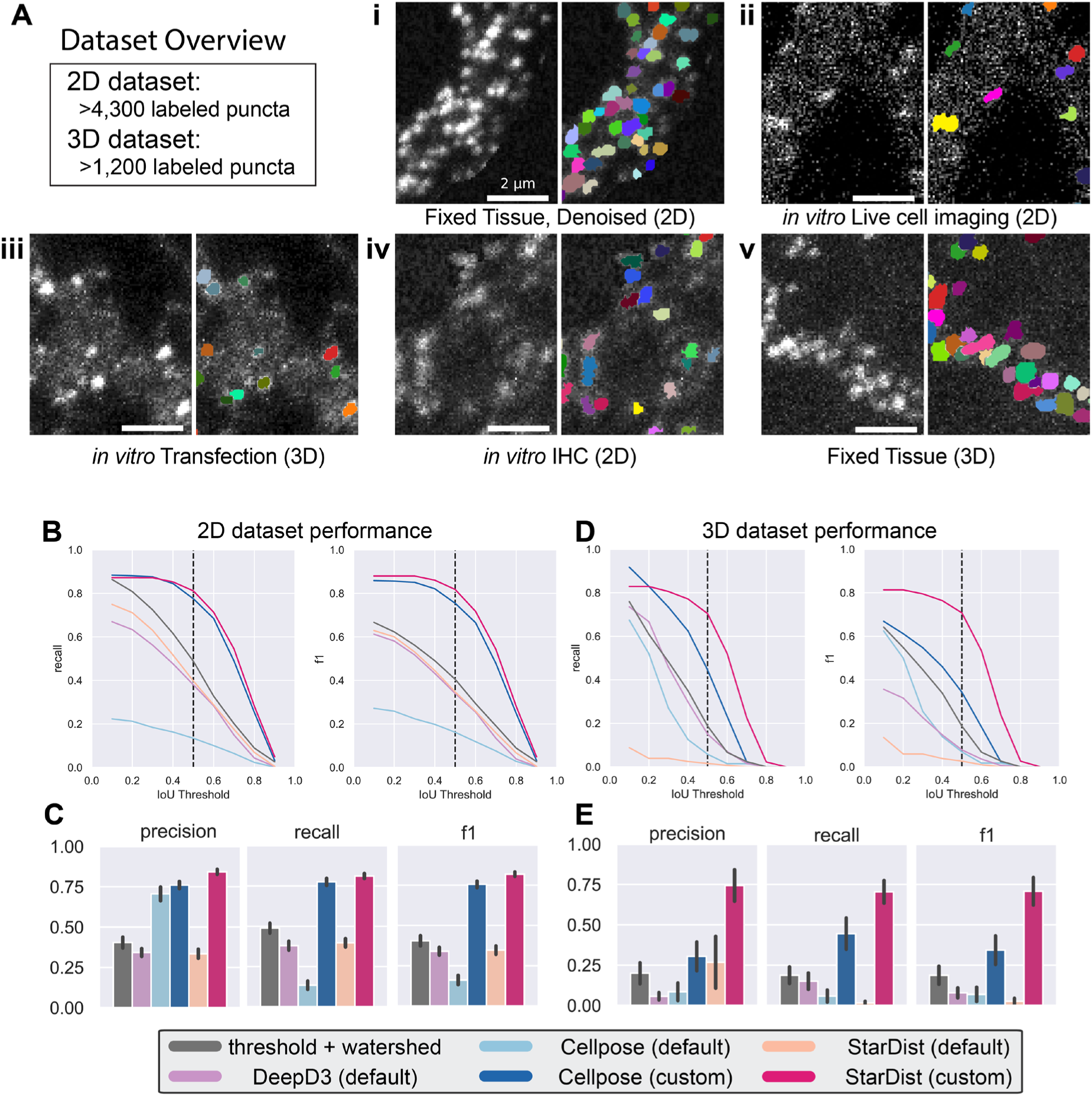
Models trained on the SynAPSeg dataset outperform pre-trained models and classical instance segmentation methods. **A)** Representative images from training dataset showing confocal maximum intensity z-projections (left half of each panel) and manually annotated instance segmentation labels (right half), with different colors reflecting unique synaptic puncta. Scale bars indicate 2µm. Panels show: **i)** PSD95-GFP labeled in fixed tissue from vGat-cre;PSD95-GFP mouse, signal was denoised using N2V ^82^; **ii)** putative inhibitory neuron from primary neuron culture expressing a PSD95-GFP FingR intrabody, image was acquired during live cell imaging; **iii)** putative excitatory neuron from fixed neuronal culture expressing PSD95-GFP FingR intrabody; **iv)** neuronal culture (fixed) where bassoon has been labeled through IHC; **v)** PSD95-GFP in tissue from vGat-cre;PSD95-GFP mouse. **iii** and **v** show images that were labeled in 3D. **B)** Evaluation of 2D segmentation methods on validation fold of the training data, recall (left) and F1 (right) scores were calculated at different IoU thresholds (top). Bar plots show the mean precision, recall, and F1 scores across validation set at IOU = 0.5 (bottom). Error bars display SEM. **C)** Same as in B but for 3D data.

These punctate structures include GABAergic synapses as well as glutamatergic synapses received by inhibitory interneurons. Their dendritic localization and dense spatial clustering can pose detection challenges that spines do not face. By including puncta from both spine and aspiny synapses, as well as presynaptic markers, our datasets capture a wide range of morphologies and expand the potential for comprehensive synapse analysis. Throughout, we use the term synaptic puncta to refer generally to fluorescently labeled clusters of synaptic proteins whether they are presynaptic, postsynaptic, and either aspiny or located at dendritic spines. To generate ground truth labels for training the models, four human experts manually annotated synaptic puncta by assigning unique pixel value identifiers to each instance. Each expert labeled different images to mitigate systemic bias. Labeling was performed either within defined regions of interest (ROIs) or on the whole field of view (**Fig S1A**).

We then used this dataset to train 2D and 3D versions of the StarDist and Cellpose-SAM models. As they have fundamental differences in their implementations, the specific parameter values were individually tuned to elicit the best performance (see **Methods**). Models were trained for a sufficient number of epochs until loss plateaued while using a learning rate that decayed as validation loss decreased (**Fig S1B-E**). We noted a persistent generalization gap in the 3D model training curves, characteristic of training on limited data. We were able to force loss convergence by using smaller training patches to increase the number of examples, but this resulted in lower segmentation accuracy on our primary benchmark dataset. Consequently, we prioritized empirical performance for our target task by utilizing the more accurate model presented here, while making the alternative regularized weights publicly available. We compared these custom trained models to pre-trained versions as well as a pre-trained deepD3 model which is designed to detect synaptic spines. To establish a baseline of performance we compared these deep learning approaches to conventional image analysis methods: thresholding followed by watershed segmentation. Since thresholding-based methods are sensitive to the parameters used, we optimized the parameters (thresholding algorithm, gaussian blur, distance threshold) via grid search for each image using the ground truth labels as targets. Each method was evaluated in 2D and 3D contexts, using the validation fold of the respective dataset. Performance was based on the intersection over union (IoU) score; we used a standard IoU threshold of 0.5 to classify detections as true positives, false positives, and false negatives and calculate recall, precision, and F1 scores. For the 2D dataset, the custom Stardist and CellPose models similarly had the highest precision, recall, and F1 scores (**Fig 1B,C**). In contrast, for the 3D dataset, the StarDist model performed best on all metrics (**Fig 1D,E**, see **Fig S2A,B** for example segmentations). A known limitation of the IoU metric is its disproportionate penalization of under-segmentation relative to over-segmentation, an effect that is particularly pronounced for small structures such as synapses. We therefore evaluated segmentation performance using shape-and localization-based metrics, including Hausdorff distance and centroid distance, which provide more balanced sensitivity^35^. The trends remained the same as observed with the IoU scores (**Fig S2C,D**). Since the custom StarDist model performed best across both 2D and 3D datasets we used these models for subsequent analyses and testing.

### Benchmarking model performance

To assess the custom StarDist models relative to human performance, we constructed three novel multi-rater benchmark datasets. These datasets consist of two 2D images and one 3D volume selected to represent visually distinct image contexts from the training distribution (**Fig 2A**; see **Methods** for further details). Importantly, these images were held out entirely from both the training and validation sets. For each image, five human experts provided independent, pixel-wise instance segmentation annotations.

**Figure 2.**
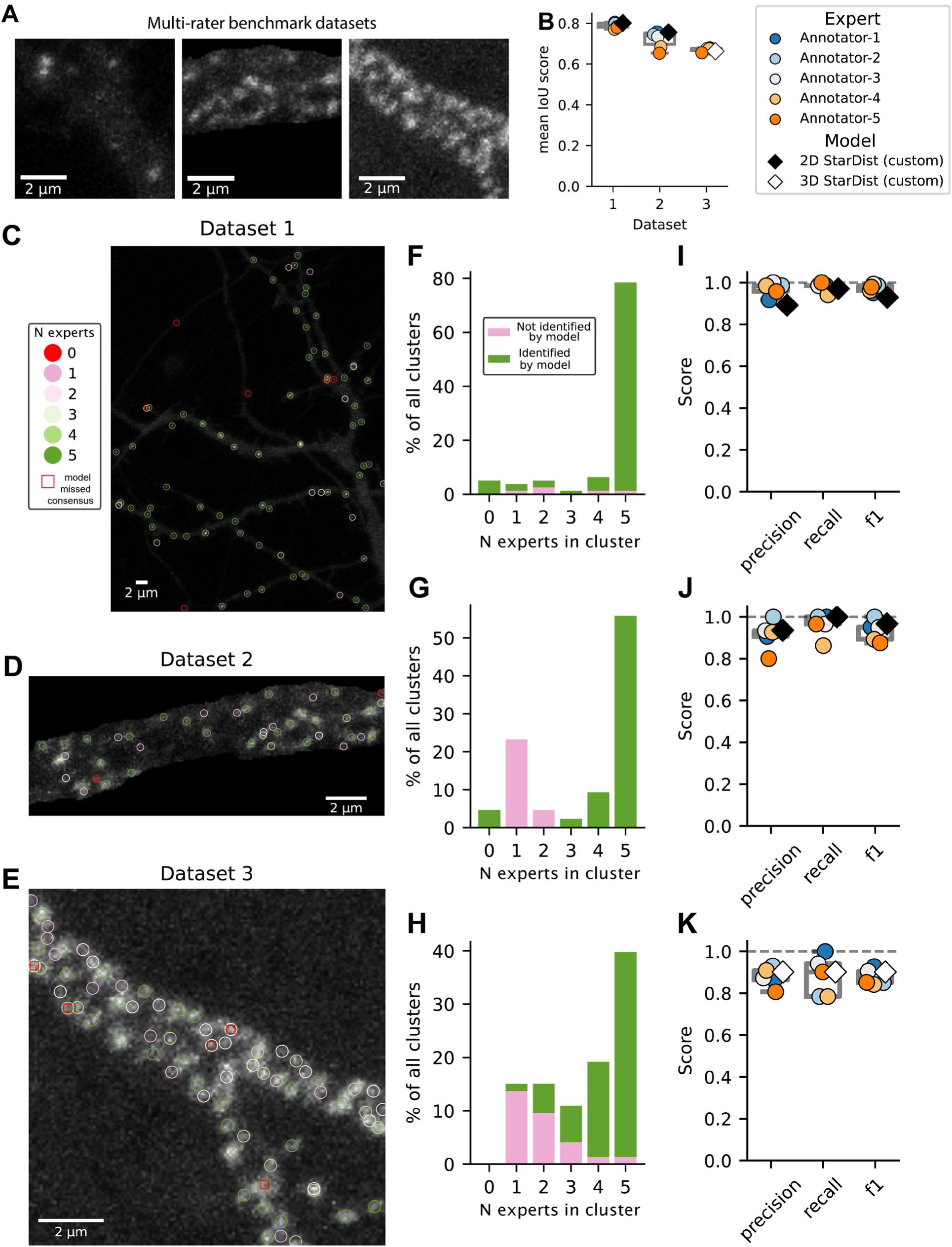
StarDist models trained on the SynAPSeg dataset reach human-level performance. **A)** Maximum intensity z-projection showing synaptic puncta in ROIs from each benchmark dataset acquired by confocal microscopy. Scale bars indicate 2μm. Dataset 1: z-projection showing *in vitro* preparation of primary neuronal culture transfected with plasmid encoding GFP-tagged PSD95-FingR. Dataset 2: z-projection showing *in vitro* preparation where Homer puncta have been labeled by immunohistochemistry. Dataset 3: volume acquired from vGat-cre:PSD95-GFP mouse brain tissue sections. **B)** Mean IoU score between each expert annotator and the StarDst models across each benchmark dataset. **C-E)** Maximum intensity z-projection showing full images of each dataset. Overlaid circles indicate expert-matched synaptic puncta aligned by spatial clustering. The circles are color-coded to reflect the number of experts in the cluster, except red circles which are detections uniquely identified by the model. Red squares indicate instances where the model did not identify a consensus cluster that the majority (n>2) experts did. **F-H)** Stacked barplots showing the percentage of experts that identified each cluster. The performance of the model relative to the human experts is indicated by the colored proportion of each bar. Green indicates the percentage of human-formed clusters that the model also identified, while pink reflects the percentage not identified by the model. The bar heights sum to 100% and reflect the total clusters identified by the experts and the model. **I-K)** Precision, accuracy, and recall scores calculated using expert consensus clusters as true positives, and false positives as clusters identified by a single expert or model. For all boxplots, the box extends from the 1st to 3rd quartiles, the center line reflects the median, and whiskers indicate full range.

Overall, the number, average intensity, and average size of model-predicted labels fell well within the range of human annotations (**Fig S3A-C**). For the 3D benchmark (Dataset 3), we noted that the model’s predictions tended to be about 20% larger in size than the median expert-annotated object, though experts showed high variability and average sizes ranging from 65 to 125 voxels. To more quantitatively evaluate human and model performance, we calculated the IoU score between each expert and the model for each image (**Fig S3D**). The mean IoU score for the models was similar to the highest scoring raters across all datasets, indicating high overlap between experts and the model (**Fig 2B**). As human annotations of synaptic puncta have been shown to have high inter-rater variability (**Fig S3E**, and see ^6^), we sought to employ evaluation metrics that capture the expert consensus as a more robust performance target. To do this, we performed spatial clustering on the centroids of labeled objects to determine the convergence of expert agreement for individual synaptic puncta (**Fig 2C-E** and see **Fig S3F** for pixel-wise segmentation masks from human annotators). This allowed us to quantify the percentage of objects the model identified relative to the number of humans that also identified that same object. Here we define consensus clusters as those identified by the majority of experts (i.e. n > 2). Overall, the model identified 95% (148/155) of consensus clusters (**Fig 2F-H**). Only 3.5% (7/195) of clusters were uniquely identified by the model—a lower rate than the percentage of clusters with a single human annotator (8.2%; 16/195). To quantify reliability, we then evaluated human-to-human against human-to-model precision, recall, and F1 scores using consensus labels as ground truth (**Fig 2I-K**). On all datasets, the model’s precision, recall, and F1 scores were comparable to human annotators. Importantly, segmentation of these datasets took up to an hour, depending on the annotator and dataset, but just seconds for the trained model. In summary, we show the custom StarDist model reaches human-level performance in a significantly shorter amount of time, highlighting the scalability of this approach.

### SynAPSeg enables fully automated deep learning-based synapse detection and quantification

To facilitate use and integration of deep learning-based tools for image analysis, we developed SynAPSeg as an open-source Python framework available across Windows, Mac, and Linux operating systems (**Fig 3**). Workflows in SynAPSeg are accessible through a graphical user interface (**Fig S4A**) and structured around three stages: segmentation, annotation, and quantification. The *Segmentation Module* provides a flexible interface for applying highly customizable deep learning-based pipelines. We include several different deep-learning models that users can chain together depending on their use case. This includes models for self-supervised denoising, like N2V2 ^36^, instance segmentation, like StarDist ^26^, Cellpose ^30^, or DeepD3 ^6^, and semantic segmentation using the Segmentation-Models library ^37^ (e.g. for dendrite segmentation). The *Annotation Module* allows users to manually correct model outputs and define custom ROIs through a Napari ^38^ interface. In addition, The *Quantification Module* provides flexible plugins for feature extraction, colocalization analysis, and ROI assignment that allow users to quantify the density and morphological features of segmented objects, regardless of what they represent (e.g. synapses, dendrites, soma). To enable integration across existing tools and platforms, SynAPSeg supports a range of image file formats through the Bio-Formats ^39^ library and utilizes an open file structure for Data Management so that external software can both read and contribute data. This interoperability allows using QuPath ^40^ and ABBA ^41^ for brain atlas registration and incorporating the region maps back into the SynAPSeg quantification pipeline. In addition, all steps generate human-readable metadata and configuration files to make analysis reproducible and transparent.

**Figure 3.**
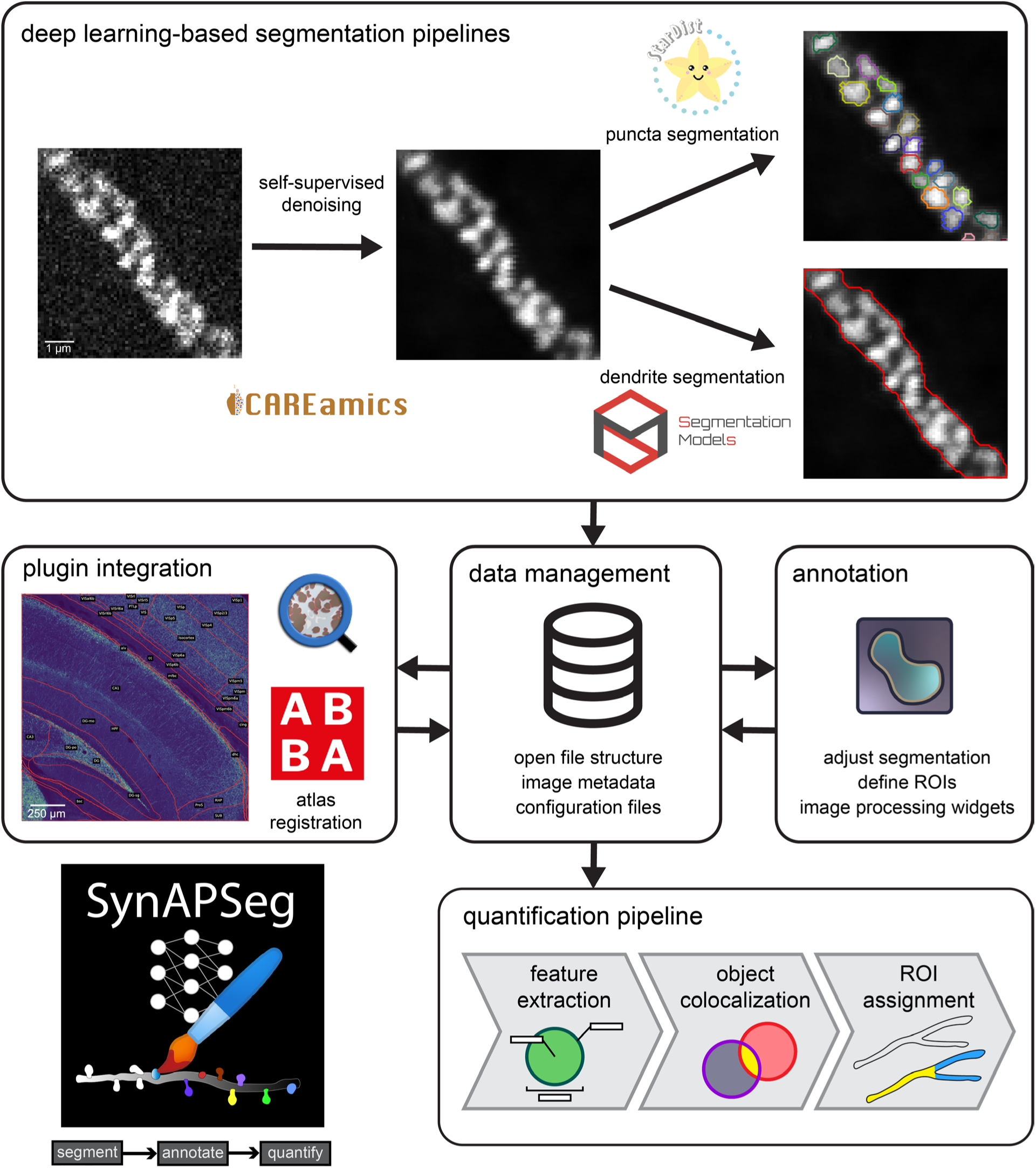
SynAPSeg provides an accessible framework for deep learning-based image analysis. The *Segmentation Module* allows users to compose flexible pipelines using state-of-the-art models for self-supervised denoising (e.g., Noise2Void via the CAREamics library) instance segmentation (e.g., StarDist). Semantic segmentation using custom models is implemented using the Segmentation Models library. Additional architectures, including Cellpose and DeepD3, are also available. *Data Management* is handled through an open file structure where images are stored as TIFF or OME-TIFF files with comprehensive metadata. SynAPSeg generates human-readable configuration files for every pipeline execution to ensure experimental reproducibility. The *Annotation Module* integrates a Napari-based GUI for manual refinement of segmentation results, definition of custom ROIs, and interactive image processing. SynAPSeg is designed for interoperability; QuPath projects can be initialized within the SynAPSeg directory structure to interface with ABBA (Aligning Big Brains & Atlases) for brain atlas registration. *The Quantification Module* provides a composable pipeline for extracting morphological and intensity features, performing multi-channel object colocalization, and assigning detected objects to semantic ROIs. All of these modules are integrated within a unified GUI, facilitating a streamlined end-to-end workflow from processing raw data to quantifying results.

To demonstrate the broad utility of the SynAPSeg platform, we analyzed 3D confocal immunofluorescence data of a primary cultured GABAergic inhibitory interneuron with pre and postsynaptic markers. Bassoon (presynaptic) and PSD95 (postsynaptic) puncta were segmented using the custom 3D StarDist model, and ROIs were manually drawn to include primary and secondary dendrites (**Fig S4B,C**). The glutamatergic synapses were largely aspiny, as expected for a GABAergic inhibitory interneuron. We performed 3D object-based colocalization analysis (see **Methods**) to identify overlapping pre and postsynaptic puncta **(Fig S4D)**. We considered co-localized puncta as putative synapses, and quantified the percentage of PSD95 that was synaptic, as well as the density of synapses within distinct dendritic compartments (**Fig S4E**). Beyond accurate density measurements, a key advantage of instance segmentation is the ability to extract features from individual puncta. This is particularly valuable given that the intensity of postsynaptic proteins, such as PSD95 and GluA1, have been shown to positively correlate with synaptic strength^1,42^. We demonstrate this analysis by extracting features of each colocalized PSD95 puncta **(Fig S4F)**. Collectively, these results demonstrate that SynAPSeg streamlines complex analytical pipelines, replacing labor-intensive manual steps with an automated workflow that enables testing fine-grained hypotheses at scale.

### Large-scale analysis of cell type-specific synaptic organization in the hippocampus

While excitatory neurons have glutamatergic synapses that typically are located in dendritic spines that can be morphologically identified, GABAergic inhibitory interneuron (INs) have glutamatergic synapses that are largely located on the dendritic shaft. This makes it more of a challenge to study glutamatergic synapse number and properties in INs. Genetic strategies, where postsynaptic glutamatergic markers are fluorescently labelled, provide a means to visualize aspiny synapses. For example, PSD95 is typically used as a glutamatergic synapse marker, and can be endogenously labelled with a transgenic strategy ^1,42,43^ or with PSD95-targeting intrabodies ^2,3,44,45^. Data from PSD95-GFP CreNABLED mice ^43^ were included in our training dataset (**Table S1, Fig 1A**).

The density of INs is known to vary by subregion, and distinct anatomical projections are known to target INs in specific hippocampal subregions. However, the relative densities and features of IN glutamatergic synapses across the hippocampus is unknown. Previous studies have evaluated how glutamatergic synapses containing PSD95 and SAP102 in all neurons are distributed in the brain, including in the hippocampus ^46^. Furthermore, they identified age-dependent changes in the synaptic architecture in aged animals ^47,48^. We sought to characterize the organization of glutamatergic synapses in INs within the hippocampus and evaluate if this changes with aging. We used a genetic strategy where endogenous PSD95 was labelled with GFP selectively in INs (vGAT-Cre;PSD95-GFP mice; see **Methods**). We then used the SynAPSeg platform to quantify the properties of IN glutamatergic PSD95 puncta in the hippocampus of early and late adulthood mice. Coronal sections from vGAT-Cre;PSD95-GFP mice aged 3 or 12 months (n=5-6) were stained with DAPI and 2D tile scan images of the dorsal hippocampus were acquired using confocal microscopy (**Fig 4A** and see **Fig S5A** for zoomed in examples). We observed large autofluorescent puncta in the pyramidal and granular cell regions in older animals which we interpreted as lipofuscin-related autofluorescence, so these regions were excluded from analyses. Synapses were segmented using the custom 2D StarDist model described above. We utilized QuPath and ABBA to manually register images to the Kim Mouse Atlas ^49^ (**Fig 4B,C**). PSD95-GFP density, area, and intensity were extracted and localized to distinct hippocampal subregions using the object detection and ROI assignment plugins from the Quantification Module in SynAPSeg (**Fig S5B**). In total, this dataset comprises over 3.8 million puncta localized to 16 distinct hippocampal subregions.

**Figure 4.**
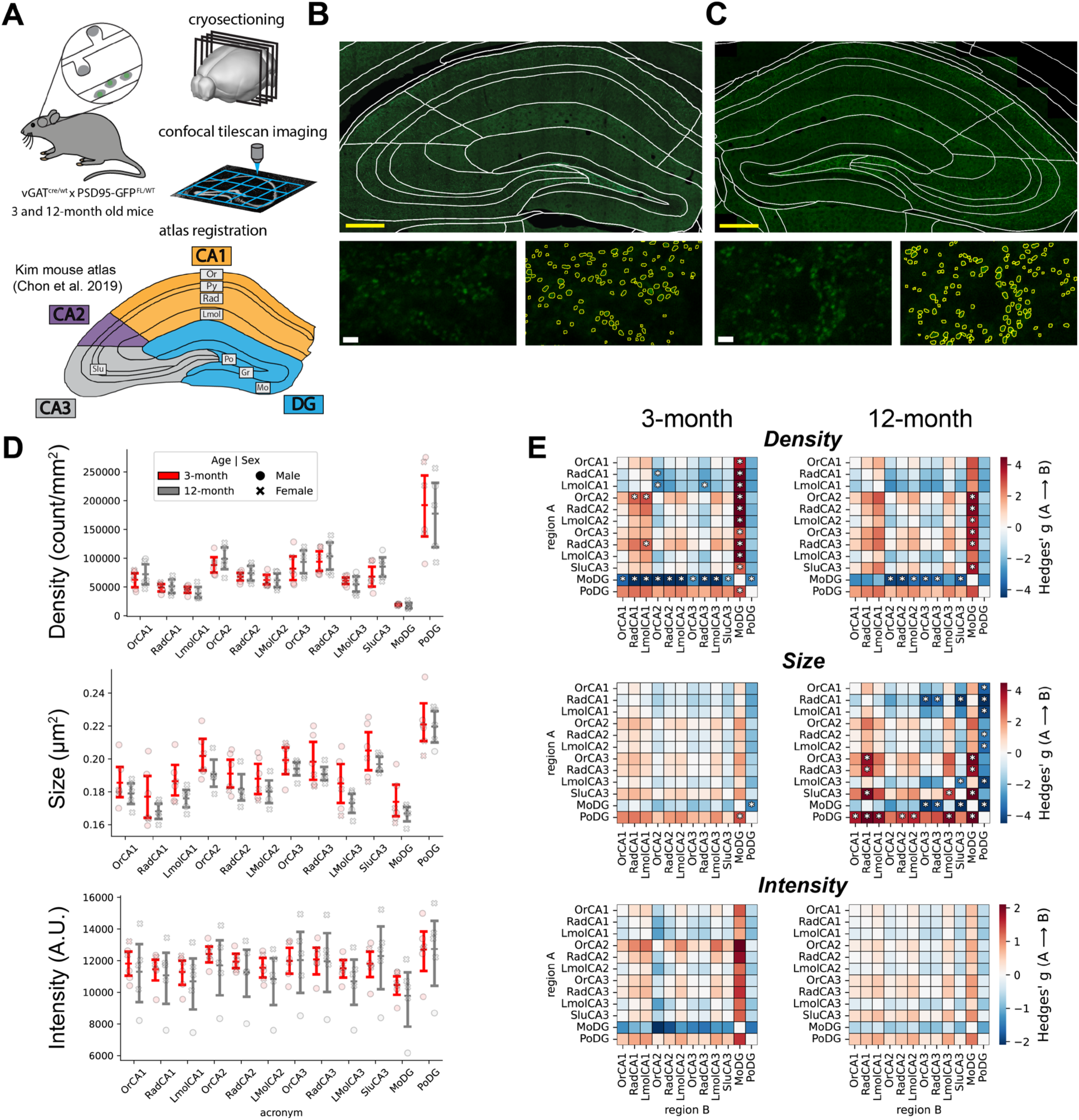
Large-scale 2D analysis of inhibitory neuron glutamatergic inputs in the dorsal hippocampus reveals conserved sub-region-specific density over adulthood **A)** Overview of approach. Coronal sections from vGAT-Cre;PSD95-GFP mice aged 3 and 12 months were imaged using a confocal microscope in tile scan mode to acquire 63x magnification images of the dorsal hippocampus. Images were registered to the Kim Mouse Atlas using ABBA. SynAPSeg was used to then segment PSD95 puncta with the custom 2D StarDist model. Puncta features (density, intensity, size) were extracted and localized to distinct hippocampal subregions using the Quantification Module. **B)** Representative image showing full FOVs acquired at 63X magnification for the 3-month group (top panel; scalebar: 250 µm), enlarged ROI showing dense glutamatergic innervation of inhibitory interneurons in the polymorphic layer of the dentate gyrus (bottom left panel; scalebar: 2 µm), and overlaid segmentation outlines (bottom right panel). **C)** same as in B but for the 12-month group. **D)** Quantitative analysis of PSD95 puncta density, mean fluorescence intensity, and puncta size across hippocampal subregions. Each data point represents an individual animal, with markers color-coded by age: 3 months (red) and 12 months (gray). Biological sex is indicated by marker style, with circles representing male subjects and’X’ symbols representing female mice. Data are summarized by error bars reflecting the mean and 95% confidence interval for each age group within each subregion. No significant age vs. region interaction was observed by two-way ANOVA (see **Table S2** for statistical details). **E)** Heatmaps show pair-wise comparisons of puncta features between regions within each age group. Values represent Hedges’ g of effect size and were calculated using the regions on the left axis relative to regions along the bottom axis. Unpaired T-tests were performed, and p-values were adjusted using Šídák’s multiple comparisons. Asterisks indicate significance based on adjusted p-values (q-values): *q < 0.05. Hippocampal subregion abbreviations are defined as follows: Field CA1-3 (CA1-CA3), Oriens layer (Or), Pyramidal layer (Py), Radiatum layer (Rad), Lacunosum moleculare (Lmol), Stratum lucidum (Slu), Dentate gyrus (DG), Granule cell layer (Gr), Molecular layer (Mo), and Polymorph layer (Po).

We observed discrete patterns of synaptic density, size, and intensity localized to specific hippocampal subregions (**Fig 4D**). Across these measurements, the poDG showed the largest and most dense puncta of all hippocampal subregions, followed by CA3 subregions.

These patterns were largely consistent across age groups. Two-way ANOVA of synaptic density indicated 72% of variance was explained by region with no significant effect from age or age x region interaction (See **Table S2** for full statistical details for this and all subsequent analyses). To identify larger trends in these properties, we then quantified inter-region differences within age groups using pair-wise multiple comparisons and are expressed as Hedges’ g of effect size (**Fig 4E**). In both age groups, we saw some significant differences between inter-regional PSD95-GFP properties. A greater number of regions showed significant changes in puncta size in the 12-month group, while the 3-month group showed slightly more inter-regional density differences. No significant trends in PSD95 intensity were observed. The vast majority of all differences occurred between the moDG and other brain regions, it notably showed the lowest absolute values across all features. Overall, these data indicate patterns of synaptic density were largely conserved with age and demonstrate the utility of SynAPSeg for fully automated segmentation and quantification of synaptic puncta in large datasets.

### Analysis of PSD95 puncta on dendrites of CA1 PV Interneurons in 3-and 12-month mice

INs represent just 10-15% of all neurons in the hippocampus but are incredibly diverse with respect to their molecular profiles, electrophysiological properties, and their roles in neuronal circuits ^50–53^. In the hippocampus, parvalbumin positive INs (PV-INs) make up roughly 25% of all INs ^54^, and are important in CA1 for the generation of gamma oscillations which have been shown to play a critical role in information processing and memory encoding ^55–58^. Aging is associated with a cognitive decline, and reductions in gamma power ^59,60^. While we did not see broad changes in the organization of glutamatergic synapses in all INs with aging **(Fig 4**), it remains possible that specific subtypes of INs are impacted with differences masked when looking at all INs. Because of their link to circuit and cognitive function, we chose to evaluate glutamatergic synapse organization specifically in PV-INs within the CA1 with aging.

To address this, coronal sections from vGAT-cre;PSD95-GFP mice aged 3-and 12-months were sectioned and PV-INs identified with IHC. Volumetric confocal images were acquired along the radiatum and oriens layers in the CA1 (**Fig 5A**). PV-IN dendrites were manually drawn using SynAPSeg’s annotation module to generate 3D ROIs. The mean intensity of PV immunoreactivity was extracted using the 3D dendrite ROIs, averaged within animals, and compared between age groups (**Fig 5B, C**). The 12-month group showed significantly lower parvalbumin intensity compared to 3-month animals (**Fig 5C**). The size and length of PV-IN dendrites analyzed were not significantly different between groups **(Fig S5A)**. Next, we used SynAPSeg to quantitatively assess changes in PV-IN glutamatergic synapse number and properties with age, using PSD95-GFP as a proxy for glutamatergic synapses. To do this we used a segmentation pipeline consisting of a N2V2 self-supervised denoising and our custom 3D StarDist model to detect PSD95-GFP puncta (**Fig 5D**). The segmented PSD95-GFP puncta were assigned to PV dendrites using the 3D quantification module in SynAPSeg. The properties of PSD95 puncta along PV dendrites were then analyzed and features including the density, area, and intensity were extracted. The PSD95-GFP linear density was positively correlated with the intensity of parvalbumin expression in both age groups (**Fig S5B**), consistent with previous reports ^61^. While the intensity (**Fig 5E**) and size (**Fig 5F**) were not significantly different, the linear density (count/μm) of PSD95-GFP puncta was significantly reduced in the 12-month group (**Fig 5G**).

**Figure 5.**
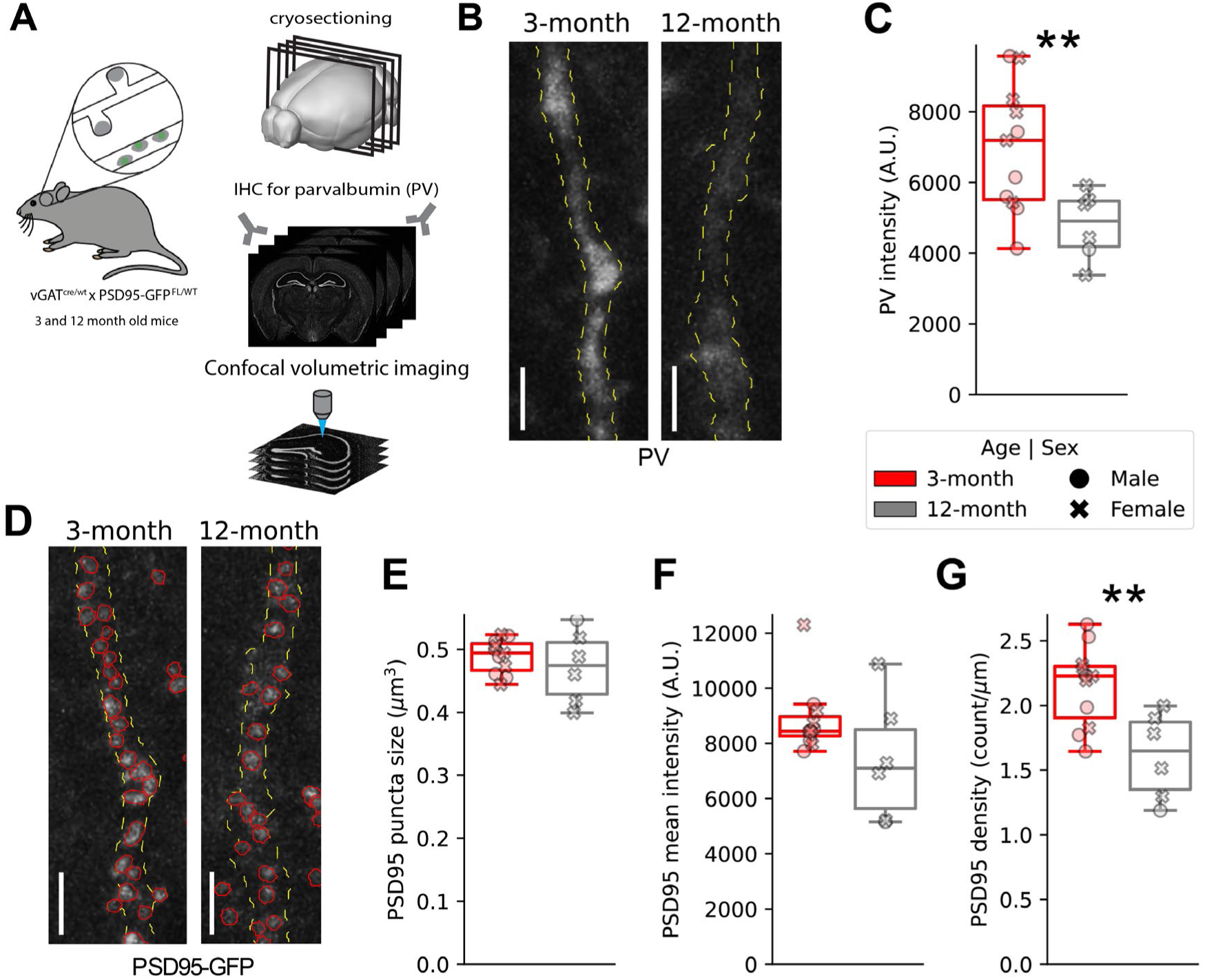
Reduced PV immunoreactivity and PSD95 density along dendrites in CA1 of aged animals. **A)** Experimental workflow: Coronal sections from 3-and 12-month-old vGAT-cre;PSD95-GFP mice were processed via IHC for PV. PV-positive dendrites were imaged in the radiatum and oriens layers of CA1 using volumetric confocal microscopy. **B)** Representative z-projections of PV-positive dendrites and manual 3D reconstructions (yellow dashed lines) in 3-month (left) and 12-month (right). Scale bars indicate 2 µm. **C)** Boxplots compare mean parvalbumin fluorescence intensity within 3D dendritic ROIs. Data points indicate the mean value of several (N = 5-11; average = 7) dendrites for each animal. PV expression is significantly reduced in 12-month animals (**p < 0.01, Welch’s T-test). **D)** Representative images of PSD95-GFP puncta along the same dendrites from B. Red outlines indicate segmented puncta detected by the custom 3D StarDist model. Scale bars: 2 µm. **E-G)** Quantitative assessment of PSD95-GFP puncta properties. **E)** puncta size (µm^3^), **F)** mean fluorescence intensity (A.U.), and **G)** linear density (count/µm). Linear density is reduced in aged animals (**p < 0.01, Welch’s T-test), while intensity and size remain stable. For all plots, circles represent males and’X’ represents females; red indicates 3-month and gray indicates 12-month animals. N = 11 3-month (N=5 females, 6 males) and 6 12-month: (5 females, 1 male).

Previous studies have demonstrated that PV-IN PSD95 puncta exhibit reduced synapse-to-synapse variability compared to those in pyramidal neurons ^42^. To investigate the spatial relationships between these clusters, we performed a nearest-neighbor analysis (**Fig S6C**). For each PSD95 puncta (“self”) we identified the three closest puncta on the same dendrite (“neighborhood”) and calculated the difference between the’self’ intensity and the’neighborhood’ average (**Fig S6D**). To account for inter-dendritic variance, intensity values were normalized within each dendrite. In both age groups, we observed a significant correlation between self and neighborhood intensities (**Fig S6E**). Notably, no correlation was observed when neighbors were randomly shuffled within a dendrite (**Fig S6F**). Taken together, these data indicate that high-intensity PSD95 puncta are spatially clustered, and that this relationship remains stable with aging.

## Discussion

Machine learning methods have revolutionized biological image segmentation, yet the development of deep learning models for synapse detection remains hindered by the scarcity of high-quality, diverse training data. Generating such datasets is notoriously time-consuming, particularly in domains like synaptic analysis where ground truth subjectivity is high even among experts. Existing resources primarily focus on dendritic spines and lack the instance-level labels necessary to resolve individual synapses in tissue preparations with dense labels ^6^. To address these gaps, we present the first comprehensive publicly available instance segmentation dataset curated fluorescent synaptic puncta. We evaluated several state-of-the-art architectures and trained custom versions on this dataset. To make these tools available to the broader community we developed SynAPSeg, a Python-based framework which integrates segmentation, annotation, and quantification modules through an accessible interactive interface. All of this is freely available through our GitHub, where we provide pre-trained models as well as the SynAPSeg training and benchmarking datasets.

Deep learning models benefit from exposure to diverse types of data during training, and as such, we generated a comprehensive training dataset containing confocal images spanning multiple imaging conditions, resolutions, synaptic morphologies, and labeling techniques.

Though we have tried to be expansive in synaptic markers used, there is a bias for postsynaptic markers, PSD95 in particular. This bias is mitigated through the incorporation of diverse training examples which span experimental conditions, labeling techniques, and sample preparations. Our view is that these factors have a greater impact on the signal characteristics than the marker identity and thus do not impede generalizability. By making these datasets publicly available we aim to empower others to further expand the diversity of open-access synaptic data, train the next generation of models, and broaden the applications of deep learning tools.

These datasets were used to evaluate whether current deep learning tools could be trained to outperform thresholding-based approaches. Custom models significantly outperformed all other methods, with StarDist architectures achieving the highest performance across both 2D and 3D datasets. The lower performance of the custom Cellpose model was unexpected given its more recent development (2025 vs. 2018). This discrepancy likely stems from a combination of merging errors and label boundary overestimation, as reflected by a marked increase in the 3D Hausdorff distance. We suspect the strategy of training 3D Cellpose models, utilizing orthogonal 2D planes to learn 3D representations, is likely ill-suited for puncta segmentation due to the compounding effect of small object size and the inherent anisotropy of this data. The low performance of the pre-trained models in this segmentation task emphasizes that context-specific training is essential to developing robust deep learning models. In a similar vein, the low performance of DeepD3 was not surprising given its training context. However, it does highlight that models trained to detect synaptic spines do not necessarily generalize to the detection of synaptic puncta. The classical thresholding-based approach, even when optimized per image, failed to capture puncta morphology and highlights limitations in conditions of variable SNR. These results underscore that robust detection of dense synaptic puncta requires deep learning models with domain-specific training.

Consistent with previous findings in spine segmentation tasks ^4,6,21^, our analysis of inter-rater reliability confirmed high variability among human experts. We find that human experts show lower agreement on 3D datasets, highlighting the difficulty of obtaining accurate instance segmentation labels in 3D contexts. Given this subjectivity, we evaluated the custom StarDist models against a human expert consensus rather than diverging ground truth signals provided by individual raters. Across all benchmark datasets, the model’s predictions showed strong agreement with the expert consensus, reaching performance levels indistinguishable from human experts. However, one of the most striking differences is the time required to generate these labels. While human expert annotations took between 10-60 minutes, depending on the dataset, the model achieved comparable results in under 10 seconds. Together this evidence suggests that properly trained deep learning approaches offer similar performance to human experts while reducing human bias and enabling large-scale synaptic analysis.

The SynAPSeg framework was developed to make advanced deep-learning models, such as those trained in this study, more accessible to the wider research community.

SynAPSeg provides a unified GUI through which segmentation and quantification pipelines can be dynamically composed. We designed this platform to be extensible; the plugin-style architecture enables developers to incorporate new models or quantification techniques with minimal code. The framework’s open data management allows external tools to easily interface with the platform, enabling users to integrate custom analytical steps into the existing quantification pipeline. Importantly, SynAPSeg enhances experimental reproducibility by automatically logging configuration parameters for every run. While we have designed this platform with synaptic analysis in mind, it supports a wide range of microscopy image formats, dimensions, and operating systems–making it a general-purpose tool for large-scale image analysis. SynAPSeg represents a significant step forward in providing end-to-end image analysis that reduces labor-intensive manual steps while maintaining the flexibility required for complex biological datasets.

To demonstrate the utility of SynAPSeg, we performed a proof-of-concept large-scale mapping of excitatory postsynaptic densities on INs across the dorsal hippocampus. Unlike pyramidal neurons, many hippocampal INs are largely aspiny (but see ^62^) which poses unique detection challenges. Overcoming these challenges is critical to studying their synaptic organization relevant for neuronal circuit function. While we did not observe broad age-dependent differences in synapse organization across the general IN population, we found consistent inter-regional differences in PSD95 density and size. This analysis encompassing millions of PSDs is only addressable through the scalable, automated quantification presented here, and enabled us to present the first large-scale mapping of IN glutamatergic synapses in the hippocampus. One of the most striking trends was the high density of PSD95 puncta in the poDG, which to the best of our knowledge has not been previously reported as most studies have focused on IN inputs in the CA1 region or more general counts of IN soma. In addition, we observed larger puncta size in the CA3 and poDG, likely reflecting Mossy Fiber inputs on INs in these regions^63^, as this aligns with known larger terminals received by CA3 pyramidal neurons from DG mossy fibers^64–66^. This synapse type is known to be plastic^67–69^ and could represent a synaptic population to focus on in studies of learning related synaptic changes. Given that PSD95 is a reliable marker of glutamatergic synapses and their strength^1,42^, these findings provide high-resolution data useful for computational models of hippocampal function.

While the total density of IN PSD95 puncta appeared unchanged with age, we speculated that specific subtypes of INs could be impacted. We focused on PV-INs, which make up around 25% of hippocampal INs in the CA1^54^, play an essential role in generating gamma oscillations ^55,70^, and have been reported to be susceptible to age-related changes ^60,71^. We replicated the previously reported decrease in PV immunoreactivity with age^60^ and confirmed that PV expression intensity correlates with the density of glutamatergic synapses^61^.

Interestingly, our analysis revealed a significant reduction in the density of PSD95 puncta in 12-month animals. This reduced synaptic input from excitatory neurons to PV-INs may indicate a diminished capacity for these cells to regulate circuit timing, potentially contributing to age-related cognitive decline before overt neuronal loss occurs. Our results contrast with a previous report ^71^, but there were several notable differences including their use of a presynaptic maker to quantify glutamatergic synapses onto PV-INs, as well as their analysis being in the cortex.

Future studies utilizing additional age points will be essential to map the exact progression of these synaptic changes. Furthermore, it would be interesting to extend these analyses to other brain regions, and in neurodegenerative mouse models where PV-IN dysfunction has been reported ^57,72,73^.

In summary, the SynAPSeg framework provides a robust, end-to-end pipeline that improves the rigor and reproducibility of synaptic analysis. While our study utilized a genetic strategy with transgenic mice, the platform is equally compatible with viral vectors, genetically encoded intrabodies, and traditional IHC. By making our custom models and training data publicly available, we provide a reliable tool for the research community. The modular, open-source nature of the framework enables it to be easily extended to other brain regions or synaptic markers, ultimately helping expand the purview of comprehensive synaptic analysis.

## Methods

### SynAPSeg dataset generation

The training dataset is comprised of 41 confocal images containing 2,400 and 1,200 manually annotated synaptic puncta in 2D and 3D, respectively. This data was generated from several different experiments and encompasses different markers as well as acquisition settings, labeling strategies, and sample preparations as detailed in **Table S1**. All images were acquired using the same confocal microscope (Zeiss LSM980) and software (Zeiss Zen). The raw intensity images were converted from Carl Zeiss Image (proprietary file format) to the tagged image file format (TIFF) to increase the accessibility of these datasets. To generate the training labels, pixel-wise manual annotation was conducted by four human experts. To ensure high fidelity, an iterative review process was employed: initial annotations were generated by a primary expert and subsequently subjected to at least two rounds of quality control review by independent annotators, ensuring that every image was verified a minimum of two times.

Individual object instances (synaptic puncta) were labeled by assigning unique integer identifiers to pixels using SynAPSeg’s Napari-based Annotation Module. Two distinct annotation strategies were utilized (**Figure S1**): “dense” annotation, where all labels within a full field of view were segmented, and “sparse” annotation, restricted to masked regions of interest, such as along individual dendritic segments.

### Training data preparation

2D images were generated by creating maximum intensity projections of confocal z-stacks. To ensure a consistent dynamic range across heterogeneous imaging conditions and minimize pixel saturation, images were normalized between the 1st and 99.9th percentile for 2D data, while 3D data was normalized between the 1st and 99.99th percentiles. For sparsely annotated images, pixels outside the annotated ROI were replaced with Gaussian noise centered on the mean intensity of the image. To standardize inputs for neural network training, images and their corresponding labels were broken up into non-overlapping patches (**Figure S1**). Based on preliminary optimization indicating that larger fields of view improved model performance, patch dimensions were maximized to utilize the available GPU memory. 2D models were trained on patches of (256, 256) pixels. For 3D models, a patch size of (32, 128, 128) was used. Patches smaller than these dimensions were zero-padded to maintain consistent input shapes. To maximize data utility, individual 2D optical sections extracted from the 3D training volumes were incorporated into the 2D training and validation sets. This greatly increased the number of training labels and the performance of the 2D models. In optimization experiments, this training approach was more effective than training on small z projections generated from contiguous optical sections. Prior to training, label masks were preprocessed to remove artifacts smaller than 5 pixels. Then dilation was applied to close small holes within labeled objects. Finally, the data was filtered to exclude patches containing less than five distinct synaptic labels. The dataset was then partitioned into training and validation folds using an 85/15 split. This split was applied to distribute the different image modalities and maximize the heterogeneity in the training and validation sets. This resulted in 586 2D patches (287 patches came from 3D slices) and 63 3D patches available for training. The same training and validation sets were utilized for all models. The complete dataset is available in both whole-volume and patched formats.

### Selection of segmentation methods for comparison

To benchmark performance, segmentation methods were selected based on two primary constraints: the capability to perform instance segmentation (separating touching objects) and the ability to handle both 2D and 3D images. We evaluated DeepD3, a framework for synaptic spine segmentation, as well as two state-of-the-art “generalist” deep learning frameworks, StarDist and Cellpose. While we compare pre-trained and custom models for the generalist architectures, we were only able to evaluate a pre-trained DeepD3 model as training custom models requires paired spine-dendrite segmentations which are not present in our dataset. Our rationale for including DeepD3 was to understand whether architectures designed and trained for spine segmentation could translate to puncta segmentation. For our evaluation of the pretrained models, we used DeepD3’s “DeepD3_32F”, Stardist’s “2D Versatile Fluo” and “3D Demo“, and the “cpsam” weights for Cellpose. To establish a baseline for performance we evaluated these models alongside an automated thresholding and watershed segmentation approach. To provide a rigorous comparison over the diverse imaging modalities, the parameters (threshold, gaussian blur, and distance threshold) for the thresholding-based approach were optimized based on the IoU score for each image and its corresponding ground truth labels. For each image, we first generated binary masks using different automated thresholding algorithms (Otsu, Yen, Li, Isodata, Triangle, Mean filters) from the scikit-image library^74^. The binary mask with the highest IoU score was then used for watershed segmentation to separate connected components. A Euclidean distance transform was applied to the binary mask, followed by Gaussian smoothing (tested sigmas: 0.25, 0.5, 1.0, 1.5, 2.0). Local maxima were then detected using a minimum distance threshold (tested distances: 1, 3, 5, 8 pixels) and utilized as seeds for the watershed transform. Artifacts smaller than five pixels were removed.

### Parameters for training the custom Stardist and Cellpose models

The StarDist and Cellpose architectures employ fundamentally distinct algorithms and different training strategies for 3D segmentation. StarDist utilizes a deep neural network to predict object geometries as star-convex polygons by estimating radial distances from a central pixel to the object’s boundary, followed by a non-maximum suppression step to resolve individual instances. For 3D StarDist models, training was performed directly on 3D volumetric patches. In contrast, Cellpose employs a “topological flow” approach where the model predicts a vector field that points toward the spatial center of each object, allowing instances to be defined by tracking these gradients to a common centroid, but limits it to using 2D slices to learn 3D representations. The recent Cellpose-SAM iteration incorporates a pre-trained Vision Transformer to enhance generalization across spatial scales and imaging conditions. For 3D segmentation, Cellpose models were trained on orthogonal 2D planes (XY, YZ, and XZ) sampled from each training volume, following the guidance outlined in the documentation.

Given the significant differences between these architectures, we selected the parameters used for training each model individually over the course of several optimization experiments. We generally found the default parameters provided by each architecture performed best. The most relevant non-default values are listed below; however, we make available the full training and hyperparameter specifications, along with the model weights (see **Data Availability**). Models were trained using learning rate scheduling that decayed as validation loss stabilized. The number of training epochs was experimentally derived based on when loss plateaued, 120 for StarDist 3D, 100 for Cellpose 3D, and 1000 epochs for the 2D models. We used learning rate of 3E-5 with decay factor of 0.5 for StarDist models. The Cellpose models used a learning rate of 5E-5 and decay of 0.1 and learning rate of 3E-5 and decay rate of 1E-5 for 2D and 3D, respectively. We employed train-time data augmentation strategies, which differed slightly by framework, and tested several different augmentation steps and parameters. We ultimately found that the simplest procedures produced the best results.

For StarDist this included random flips, rotations, and Gaussian noise which was implemented using the Python libraries Albumentations ^75^ for 2D and Volumentations ^37^ for 3D augmentation. Since external augmentation pipelines are not supported in Cellpose’s high-level API, we utilized their default internal augmentations except for scaling, thus the addition of random Gaussian noise was not applied for the Cellpose models. Models were trained using a Nvidia RTX 4090 graphics card on Windows in Conda environment running python 3.10.16, stardist 0.8.4, TensorFlow 2.10.1, cellpose 4.0.6, and PyTorch 2.9.1 (full environment builds for windows, linux, and mac operating systems can be found on our GitHub repository).

### Evaluation of segmentation methods

To quantify the performance of these different algorithms we evaluated segmentation accuracy using standard computer vision metrics, including Intersection over Union (IoU), accuracy, recall, and F1 scores. All methods were evaluated independently on the validation folds of the 2D and 3D datasets. The IoU score was utilized as the primary metric for comparison due to its prevalence as a standard benchmark in the machine learning field, despite recognized limitations regarding small-object segmentation ^6^. Predicted synaptic puncta were classified as True Positives (TP), False Positives (FP), or False Negatives (FN) based on the IoU overlap with ground truth masks. We calculated mean recall and F1 scores across a range of IoU thresholds (**Fig 1B,D**). Objects with an IoU score > 0.5 were considered TPs for the purpose of selecting the best-performing model (**Fig 1C, E**). To provide a complementary assessment of boundary precision and localization accuracy, we also calculated the Hausdorff distance and centroid distance (**Fig S2C,D**), which offer more egalitarian penalization of over-and under-segmentation compared to overlap-based metrics ^35^. Both metrics were applied on matched pairs. Hausdorff distance was calculated using Scipy’s implementation ^76^. Centroid distance was calculated as the Euclidean distance between the centroids of the ground truth label and nearest matched predicted label.

### Benchmark dataset generation and evaluation

The multi-rater benchmark dataset consists of three distinct images (two 2D images and a 3D volume), selected to represent the diversity of imaging modalities present in the broader dataset (see **Table S1** for detailed description of different modalities). Critically, these images were held out from the training set. Each image was manually annotated by five experts using the protocols described above. We chose to only evaluate the custom StarDist models against these multi-rater benchmark datasets, as this architecture yielded the highest performance in the initial validation.

We quantified model performance using two approaches. First, we evaluated pixel wise similarity by calculating the mean IoU for every unique pair of human raters and compared these to the human-model IoU scores to determine if the model fell within the range of human variability. While this addresses pixel level agreement, utilizing a varying target as ground truth is problematic, as others have described. Therefore, we also utilized a distance-based clustering approach to establish a human expert-defined “consensus” that accounts for subjectivity in synaptic puncta identification. This was implemented following the approach as previously described ^6^. Centroids of annotated objects from all experts were extracted and spatially overlapping annotations were identified using DBSCAN to cluster annotations within a proximity threshold of three pixels in all dimensions. If this resulted in a cluster containing multiple annotations from the same expert, we then used K-means to separate it into distinct clusters. Consensus objects were defined as clusters identified by a majority of experts (n > 2). We stratified individual annotations into Confusion Matrix Components: TPs were assigned detections that identified a consensus cluster, FNs were assigned when a consensus cluster was missed, and FPs were clusters identified by a lone annotator (whether this was a human expert or the model). These components were then used to calculate classification performance metrics (accuracy, recall, and F1 scores) and jointly quantify model and expert human performance. It is important to emphasize this strategy inflates the performance of human experts as the models have no influence in determining what constitutes consensus. The confusion matrix components were also used for measuring inter-rater reliability, which was assessed through Cohen’s Kappa score using the scikit-learn libraries implementation ^77^.

### SynAPSeg Design

The SynAPSeg framework is structured around three steps: segmentation, annotation, and quantification. This core workflow is accessible through a graphical user interface that facilitates batch processing by allowing the user to define a set of parameters which will be automatically applied to all images in the project and allows settings to be applied to other projects with minimal adjustments. *Segmentation.* To initiate the pipeline, users generate a configuration file that defines data input paths, pre-processing steps, and the specific deep learning models to be deployed. This process yields an “Example” folder for each source image, which serves as a self-contained data package storing: a copy of the raw image data, the generated segmentation masks, and a metadata file recording the exact segmentation parameters and metadata extracted from the source image. In the initial segmentation stage, raw microscopy files are read using the AICSImageIO^78^ and Bio-Formats^39^ libraries, ensuring compatibility with a wide range of proprietary and open-standard imaging formats. *Annotation*.

The annotation module serves as a data quality control step, providing a Napari window to visualize the raw data and segmentation results. The module incorporates interactive widgets and annotation tools that allow users to refine the segmentation results by applying thresholding or by drawing directly on the image. These tools can also be used to define custom ROIs to constrain or categorize detected objects in subsequent quantification steps. Any adjustments to the segmentation masks or generated ROIs can then be exported and saved within the example folder. *Quantification.* The quantification module allows users to define custom pipelines for feature extraction, object colocalization across image channels, and ROI handling. The pipeline outputs data at two levels of granularity to accommodate different analytical needs. Low-level object data is stored in comprehensive spreadsheets containing extracted features, ROI assignments, and colocalization metrics for individual detected objects and ROIs. A high-level spreadsheet integrates these low-level sources with user-defined grouping variables (e.g., sample ID, treatment group, etc.) to generate a summary of the object level data for each initial source image. This summary is formatted for immediate integration into external graphing software.

While the GUI supports these core features, the underlying library contains several advanced modules that are currently accessible only through standalone Python scripts. These include specialized analysis tools for unsupervised object feature clustering and nearest-neighbor spatial analysis. Although the GUI does not currently support training custom deep-learning models (except for the Careamics denoising suite), we provide comprehensive training scripts via our GitHub repository. We also direct users toward dedicated platforms such as BiaPy^79^ and ZeroCostDL4Mic^80^ which offer accessible interfaces for custom model development. As a framework, SynAPSeg is designed to be extensible—each segmentation model and quantification stage is implemented using a plugin-style architecture making it simple to incorporate new features. Any externally trained model supported by a SynAPSeg plugin can be integrated into the segmentation pipeline.

### Animals used and tissue preparation

All experiments were approved by Tufts IACUC protocol. To express GFP-tagged PSD95 in inhibitory neurons, Psd95-GFP CreNABLED mice ^1,42,43^ were crossed with vGat-Cre mice (Jax #028862). All mice used in the aging experiments (**Figs 4 and 5**) were heterozygous for both alleles. Genotyping was performed by TransnetYX. Preparation of mouse brain tissue was performed uniformly across dataset generation and aging experiments. Animals were deeply anesthetized using intraperitoneal injection of pentobarbital and transcardially perfused with 35ml of ice-cold 1X phosphate-buffered saline (PBS) followed by fixative solution (4% PFA (Electron Microscopy Solutions), in PBS, pH 7.4). Brains were extracted and post-fixed for 2 hours in fixative solution at 4°C, rinsed in PBS, then put in 30% sucrose at 4°C until sunk.

Brains were then transferred to Tissue-Tek® O.C.T. Compound (Sakura #4583) and flash frozen in a dry ice-ethanol bath and stored at-80°C. Floating coronal sections were acquired using a Leica CM1900 cryostat. All tissue sections used in aging experiments were sectioned at 25 µm. Varying thickness (20-40µm) were acquired for generation of the training data. Sections were stored in cryoprotectant solution (1% polyvinyl-pyrrolidone (Millipore Sigma #PVP-40), 30% sucrose, 30% ethylene glycol (Millipore Sigma #102466, 1X PBS) at-20°C.

### Immunohistochemistry

Various IF/IHC protocols were utilized to generate the training dataset depending on the antibodies used (see **Table S1**). All immunohistochemistry was performed on free-floating sections. All incubation and wash steps were conducted at room temperature under constant agitation unless otherwise specified. For tissue used in **Fig 5**, sections were first washed three times for 10 minutes in 1X PBS, followed by 2 hours in blocking solution (10% normal goat serum (Vector Labs #S-1000-20), 0.3% Triton X-100, 1X PBS). Tissue was then incubated with rabbit anti-parvalbumin primary antibody (1:1000, Swant #PV27a) diluted in 1X PBS containing 5% normal goat serum and 0.15% Triton X-100 for 24 hours at 4°C. Following primary incubation, sections were washed and incubated with goat anti-rabbit Alexa Fluor 594 secondary antibody (1:1000, ThermoFisher #A-11012) in 1X PBS containing 5% normal goat serum and 0.15% Triton X-100 for 2 hours. After a final wash series, sections were placed in 0.1 M phosphate buffer for mounting onto SuperFrost Plus glass slides (Fisher Scientific #22-037-246). Once excess moisture had evaporated, sections were coverslipped (Millipore Sigma #CLS2980245) using 200 µL of ProLong Glass Antifade Mountant (ThermoFisher #P36984).

Sections used in **Fig 4** were washed, stained with DAPI (1:5000; ThermoFisher #62248) for 10 minutes, and washed again prior to mounting.

### Culture preparation and immunocytochemistry

Timed pregnant Sprague-Dawley rats were purchased (Charles River) and dissected at embryonic day 18. Hippocampi of 4-5 embryos were dissected into ice-cold dissection media (10x HBSS (Gibco), Penicillin-Streptomycin (Gibco), sodium pyruvate (Gibco), 10mM HEPES (Gibco), 30mM Glucose (Sigma) and Milli-Q water) prior to chemical dissociation via papain (Worthington) and subsequent trituration via P1000 pipette. Cells were plated at 100,000/well onto glass coverslips coated with Poly-L-Lysine (Sigma) of a 12-well plate (for immunofluorescence). Neurons were grown in Neurobasal media (Gibco) supplemented with Penicillin-Streptomycin (Gibco), 2mM Glutamax (Gibco), 5% horse serum (Gibco), and 2% B27 (Gibco). A full media change was performed two hours after plating cells. One day after plating cells (DIV1), 70% of media was replaced with serum-free Neurobasal media. Cells were fed once a week and maintained at 37C with 5% CO2. To express plasmids, neurons were transfected at DIV12 with Lipofectamine 2000 (ThermoFisher #11668027) mixed with plasmid DNA for 45 minutes followed by a full media change containing 50% media retained prior to transfection and 50% freshly prepared Neurobasal supplemented with B27.

### Immunocytochemistry

Prior to fixation, cells were washed once with sterile room temperature PBS. Cells were then fixed with 4% paraformaldehyde and 4% sucrose in PBS for 15 minutes at room temperature then washed three times with PBS. Prior to antibody labeling, fixed cells were permeabilized with 0.2% Triton-X in PBS for 20 minutes, then washed 3 times with PBS before blocking in 10% Normal Goat Serum and 5% Bovine Serum Albumin (Sigma-Aldritch) in PBS for 1 hour. Primary antibodies were added in the same blocking buffer and left overnight at 4°C. The next day, cells were washed 3 times with PBS before adding secondaries for 1.5 hours at room temperature, in the same buffer as primary and blocking. See **Table S1** for a list of antibodies and concentrations used in this study. Following secondary incubation, cells were washed three times in PBS followed by a single rinse in distilled water prior to mounting on SuperFrost Plus glass slides with ProLong Glass Antifade mounting media. Slides were maintained in the dark at 4°C. For **Fig S4**, the following antibodies were used: FluoTag®-X2 abberior STAR 635P anti-PSD95 (1:5000; NanoTag Biotechnologies #N3702-Ab635P-L), guinea pig anti-bassoon (1:1000; Synaptic Systems #141318), mouse IgG2A anti-GAD67 (1:500; Millipore #MAB5406), DyLight 405 anti-guinea pig (Jackson Immuno Research #106-475-003), Alexa Fluor 488 anti-mouse IgG2A (1:1000; ThermoFisher #A21131).

### Confocal imaging and data analysis using SynAPSeg

All imaging was performed on a Zeiss LSM 980 confocal microscope using a 63× objective and Zen Software (Zeiss). The confocal image used in **Fig S4** was acquired using a pixel spacing of 0.500 x 0.071 x 0.071 µm (Z, Y, X dimensions), sequential laser scans, and excitation wavelengths of 353, 493 and 653 nm. Analysis of was performed in SynAPSeg.

Basson and X2-PSD95 puncta were segmented using the custom StarDist 3D model. A small GAD67-positive segment of primary and secondary dendrite was manually annotated to generate 3D ROIs. For quantification, puncta size, intensity, and density were extracted and assigned to primary or secondary dendrites using the ‘Overlap3D’ ROI assignment algorithm. Colocalization was performed on the bassoon and PSD95 puncta using an IoU threshold of 0 so that if objects overlapped by one voxel, they would be considered putative synapses. The percentage of colocalized PSD95 was calculated by dividing the number of bassoon-positive PSD95 puncta by the total number of PSD95 puncta. The linear density (puncta/µm) was quantified by dividing the number of PSD95 puncta on each dendrite segment by the dendrite’s length.

Imaging and analysis of tissue sections from vGAT-Cre:PSD95-GFP mice was performed blind to age group (3-month or 12-month). For the experiments in **Fig 4**, each image was acquired over the entire dorsal hippocampus on one hemisphere using Zen’s tile scan mode and pixel size of 0.071 µm. A rectangular ROI, consisting of several tiles, was drawn over the imaging region and focus points were marked at the brightest Z position at each corner to acquire a single optical section over the entire imaging region. This was done to accommodate any unevenness in the tissue from mounting. DAPI and endogenous PSD95-GFP signal were acquired using sequential scans, 2x averaging, and excitation wavelengths of 353 and 493 nm. The laser intensity and gain for PSD95-GFP was optimized on control tissue. Images were excluded if there was Z-axis drift. 2-3 images were acquired for each animal. Acquired images ranged in size from 14514 to 46110 pixels along any one dimension, with median dimensions of 20352 x 33230 pixels (Y, X).

For the tile scan experiments in **Fig 4**, Carl Zeiss Image files were segmented in SynAPSeg using the 2D StarDist model to detect PSD95 puncta. The “output_image_pyramid” setting was checked to save the intensity images as multi-resolution image pyramids for compatibility with ABBA and a QuPath project was created for each image. Registrations were then performed in ABBA using the “kim_mouse_10um” atlas which contains brain region annotations at a finer parcellation level (e.g. oriens layer of CA1) than is represented in the most commonly used Allen CCFv3. For each brain section we performed a manual affine “interactive transform” to roughly position the slice, followed by a manual spline registration to position the finer structures using landmarks such as the granule cell, pyramidal, and slm layers. The atlas registrations were exported as.geojson files using a custom QuPath groovy script (available on our GitHub) and processed in SynAPSeg along with the puncta segmentation masks and intensity image using the ABBA Quantification Module. Each.geojson file stores the atlas regions as collections of polygons that have been deformed to the images. To assign objects to particular brain regions, the centroid of each puncta is used to determine the regions that contain it using an optimized implementation of the ray casting algorithm. The density, size, and intensity of PSD95 puncta in each brain region was aggregated across sections from the same animal by weighted averaging. Inter-regional differences were calculated within age groups using Hedges’ g of effect size. Multiple paired T-tests using Sidak’s correction were performed to evaluate significance.

To generate the images used in **Fig 5**, we imaged the hippocampal CA1 region of tissue from vGAT-Cre:PSD95-GFP mice that had been labeled for PV as described above. One confocal z-stack (23 x 2639 x 2639 voxels) with voxel size of 0.460 x 0.085 x 0.085 µm (Z, Y, X) was acquired for each animal. PV (AF-647) and endogenous PSD95 (GFP) signal were acquired using sequential scans, 2x averaging, and excitation wavelengths of 653 and 493 nm.

For the 3D analysis of PSD95 puncta along PV dendrites, a SynAPSeg segmentation pipeline consisting of the following steps was applied to each image: first a Careamics N2V2 denoising model ^36^ was trained on a randomly cropped image patch (ZYX dim size: 16, 1024, 1024) for 25 epochs using a patch size of (16, 64, 64) and otherwise default parameters (careamics v0.0.11). After training, the model performed inference over the whole volume and the resulting denoised image was processed by the custom StarDist 3D model to segment puncta. A separate N2V2 model was trained for each image to ensure optimal performance across the whole dataset and improve the segmentation of the dimmer PSD95 signal in the older samples. PV-positive primary dendrites were then manually annotated using SynAPSeg’s Annotation Module to generate 3D ROI masks. Each segment was demarcated using a unique pixel value. For quantification of PV dendrites, the size, intensity, and dendrite length features were extracted. For PSD95 quantification, puncta were first localized to individual dendrites using the ‘Overlap3D’ ROI assignment algorithm, then size and intensity features were extracted. The intensity was calculated using the raw source image, and not the denoised image. The linear density (puncta/µm) was quantified by dividing the number of PSD95 puncta on each dendrite segment by the dendrite’s length. The density, size, and intensity of PSD95 puncta was averaged over several dendrites per animal. For the nearest neighbor analysis in **Fig S6**, we examined how the intensity of each punctum (“self”) compared to its three nearest neighbors (“neighborhood”) along each dendrite. The puncta intensity values were z score normalized within each dendrite. The “neighborhood delta” was calculated as averaged neighborhood intensity minus self intensity. For the correlation analysis, we compared the self intensity to the mean intensity of its neighbors and relative to a permuted control where neighbors were randomly drawn from the pool of all puncta on that dendrite.

### Software

SynAPSeg is developed as a community tool. All code is open-source and available at our GitHub repository https://github.com/pascalschamber/SynAPSeg which also contains documentation and several demos. The code uses several open-source Python libraries, which are cited throughout.

## Data Availability

The training datasets, benchmark datasets, and the custom trained models have deposited at the following DOI: https://doi.org/10.5281/zenodo.18988899. Other datasets generated during this study and any additional information needed to reproduce them are available upon request from the lead contact.

## Statistical analysis

All statistical analysis was performed using Python Scipy and Pingouin ^81^ libraries.

Figures were generated using Python Matplotlib and Seaborn libraries. Custom scripts used to analyze data and generate figures are available on our GitHub. N represents the number of animals used in each experiment, unless otherwise noted. P < 0.05 was considered statistically significant and is presented as: *p < 0.05, **p < 0.01, ***p < 0.001. Details of all statistical tests (n numbers, age, and sex of animals) can be found in **Table S2**.

## Acknowledgements

This work was supported by funding from the National Institutes of Health (R00MH124920 to AMB, RF1MH120119 and RF1MH130784 to HZ). We are grateful for support from the Tufts University CMS staff for help with mouse breeding and husbandry.

## Supplementary Figures

**Supplemental Figure 1.**
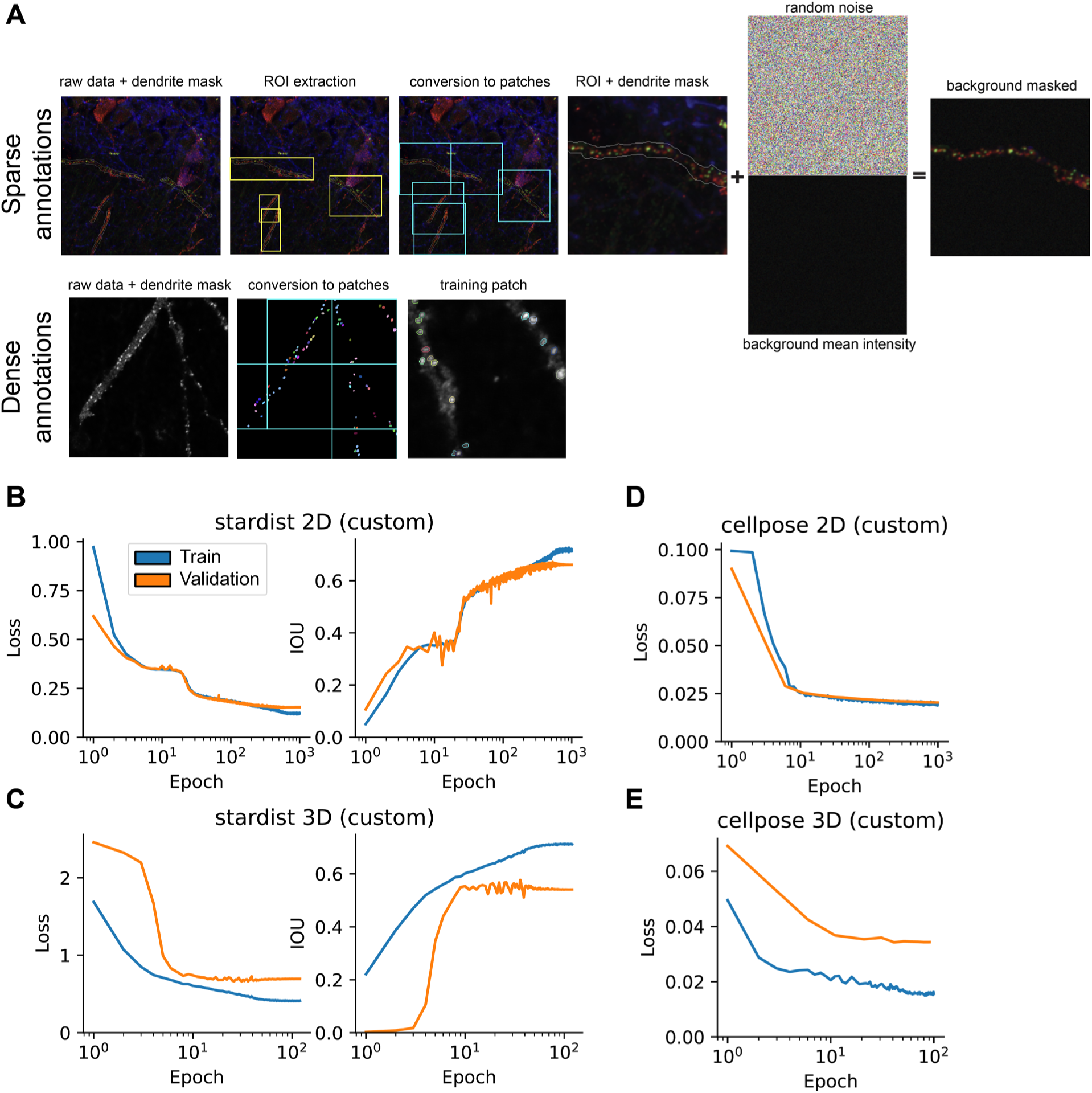
Dataset preparation and model training. **A)** Two approaches were used to convert annotated images into patches for training the models. Sparse annotations (upper panels) refer to images where only puncta within a region of interest (ROI) were labeled. Each ROI was processed independently, first masking the field of view outside the ROI, before splitting up into non-overlapping patches. All pixels outside of the ROI mask were filled with random noise drawn from a distribution centered on the mean intensity of the patch. For densely annotated images (lower panels), the entire field of view was annotated, and non-overlapping patches were extracted using a regularly spaced grid. Before training, images were processed to fill small holes and patches containing less than 5 labels were discarded. **B)** Combined training and validation loss (left panel) and IoU (right panel) during training of the 2D StarDist model on the SynAPSeg dataset. **C)** Same as in B but for the 3D StarDist model. **D)** Training loss for the 2D Cellpose model. **E)** Same as in D but for the 3D Cellpose model. Note that the models use different loss functions and values are not directly comparable. Additionally, the Cellpose library’s training module does not calculate IoU metrics and thus are not shown.

**Supplemental Figure 2.**
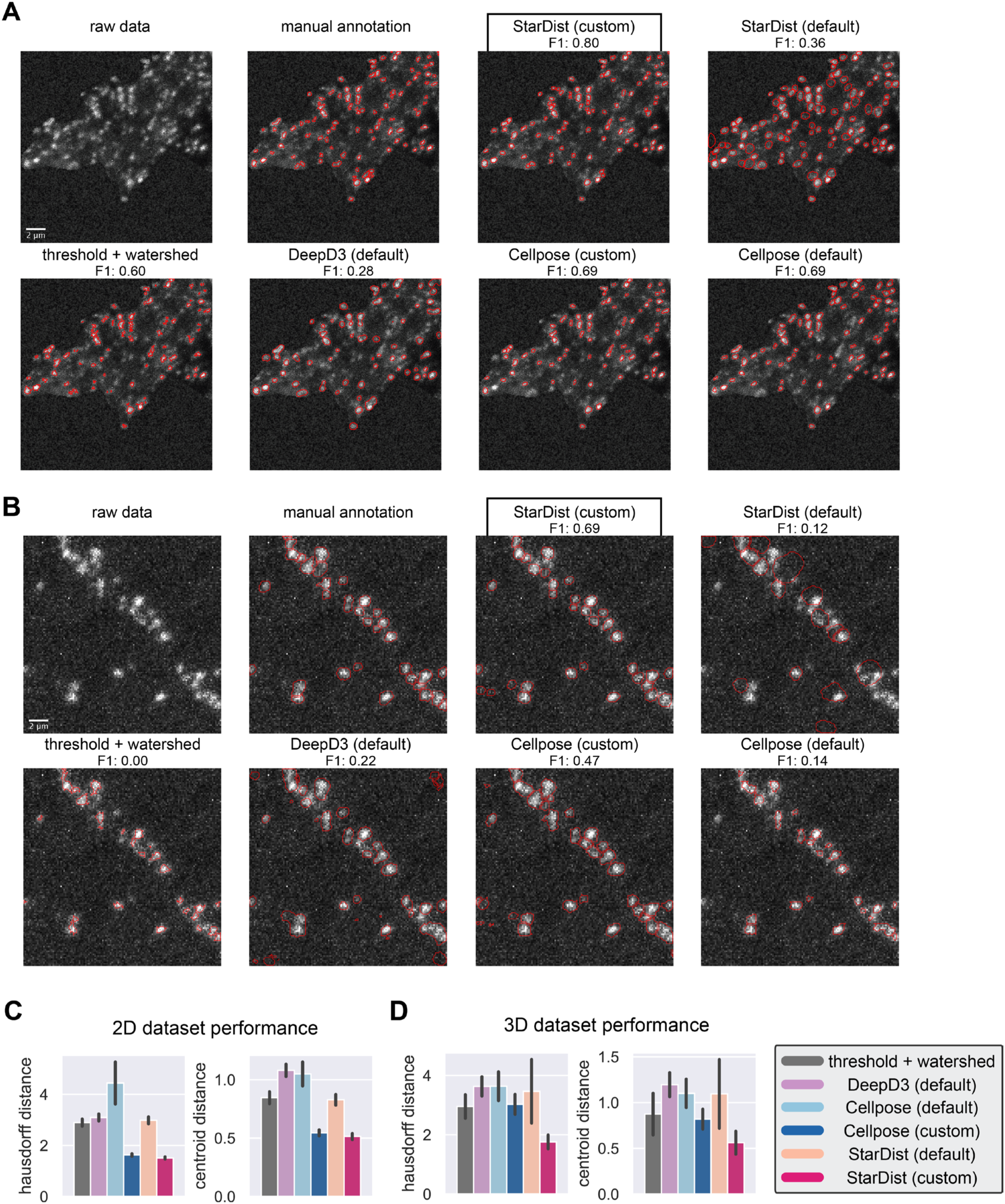
Extended comparison of instance segmentation techniques. **A)** Representative images selected from the 2D validation dataset comparing segmentation results of the different methods. Black box denotes the result with the highest F1 score. Scale bars indicate 2µm. **B)** Same as in A but for the 3D dataset. **C)** Shape/boundary (left) and centroid-based (right) metrics computed over the 2D validation dataset. Lower values indicate higher spatial precision. Hausdorff distance was calculated by comparing the distance between the sets of foreground pixels for each matched pair of ground truth and predicted instances, defined as the maximum of the shortest distances between any two points in the respective sets. Centroid distance was calculated as the Euclidean distance between the center of mass of the predicted mask and its matched ground truth. Error bars display SEM. **D)** Same as in C but for the 3D dataset.

**Supplemental Figure 3.**
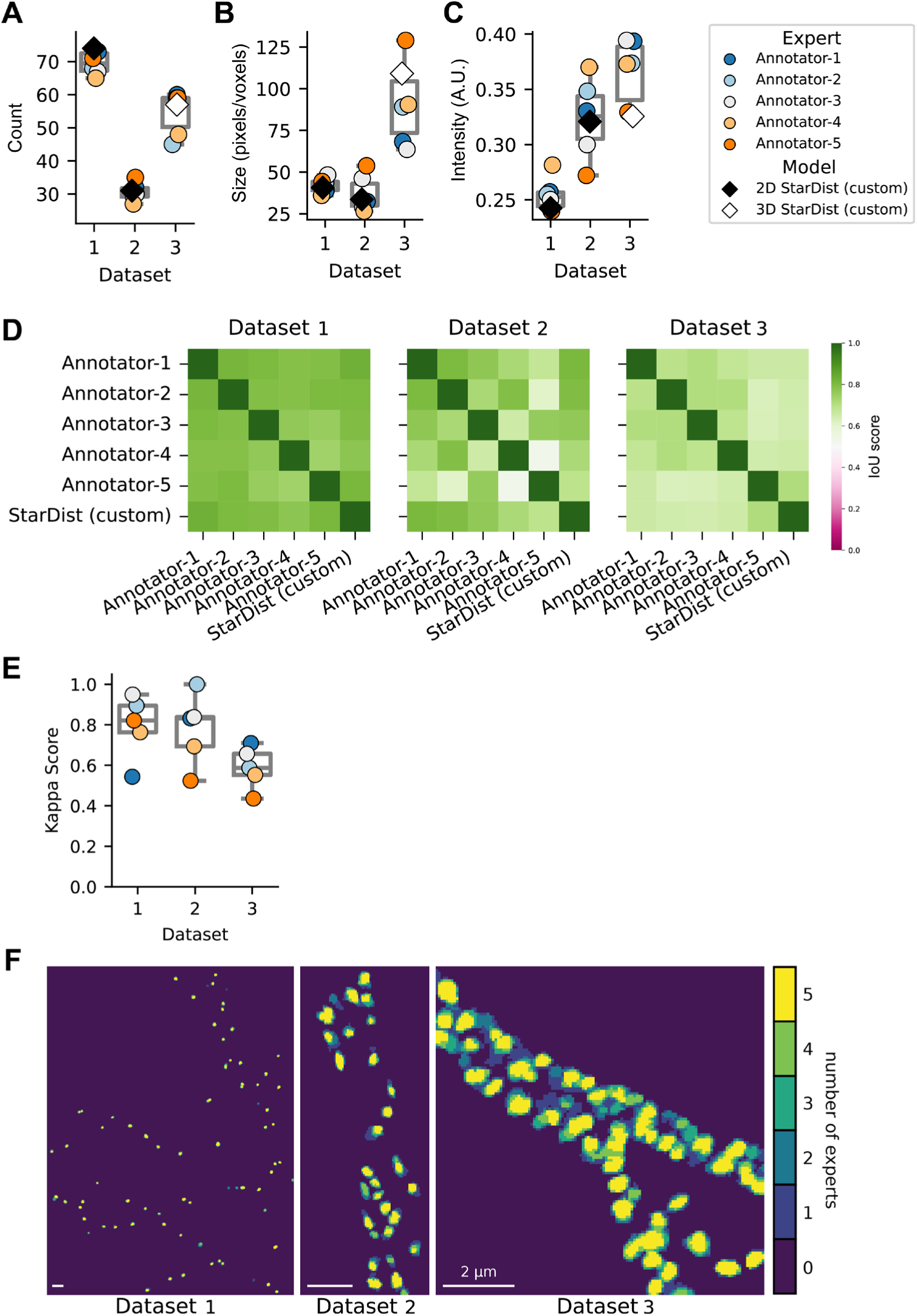
Extended benchmarking of expert annotations and performance of the custom trained StarDist models. **A-C)** Boxplots comparing morphological and intensity features of segmented objects identified by experts and the StarDist model. Features include **A)** total puncta count, **B)** mean size (in pixels for datasets 1 and 2; voxels for dataset 3), **C)** and mean fluorescence intensity. The 2D and 3D StarDist models are indicated by filled and unfilled diamonds, respectively. **D)** Heatmaps show IoU score computed between each of the five expert annotators and the custom-trained StarDist model across the three benchmark datasets. **E)** Inter-rater reliability was determined using the experts’ consensus clusters as ground truth and is measured by Cohen’s Kappa score. Each datapoint reflects the score between each of the expert annotations and the consensus labels. Values range from-1 (complete disagreement) to +1 (perfect agreement), while 0 indicates agreement no better than random chance. **F)** Images show pixel-wise segmentation masks for each of the human expert annotations that have been converted to binary pixel maps and summed to reflect the consensus.

**Supplemental Figure 4.**
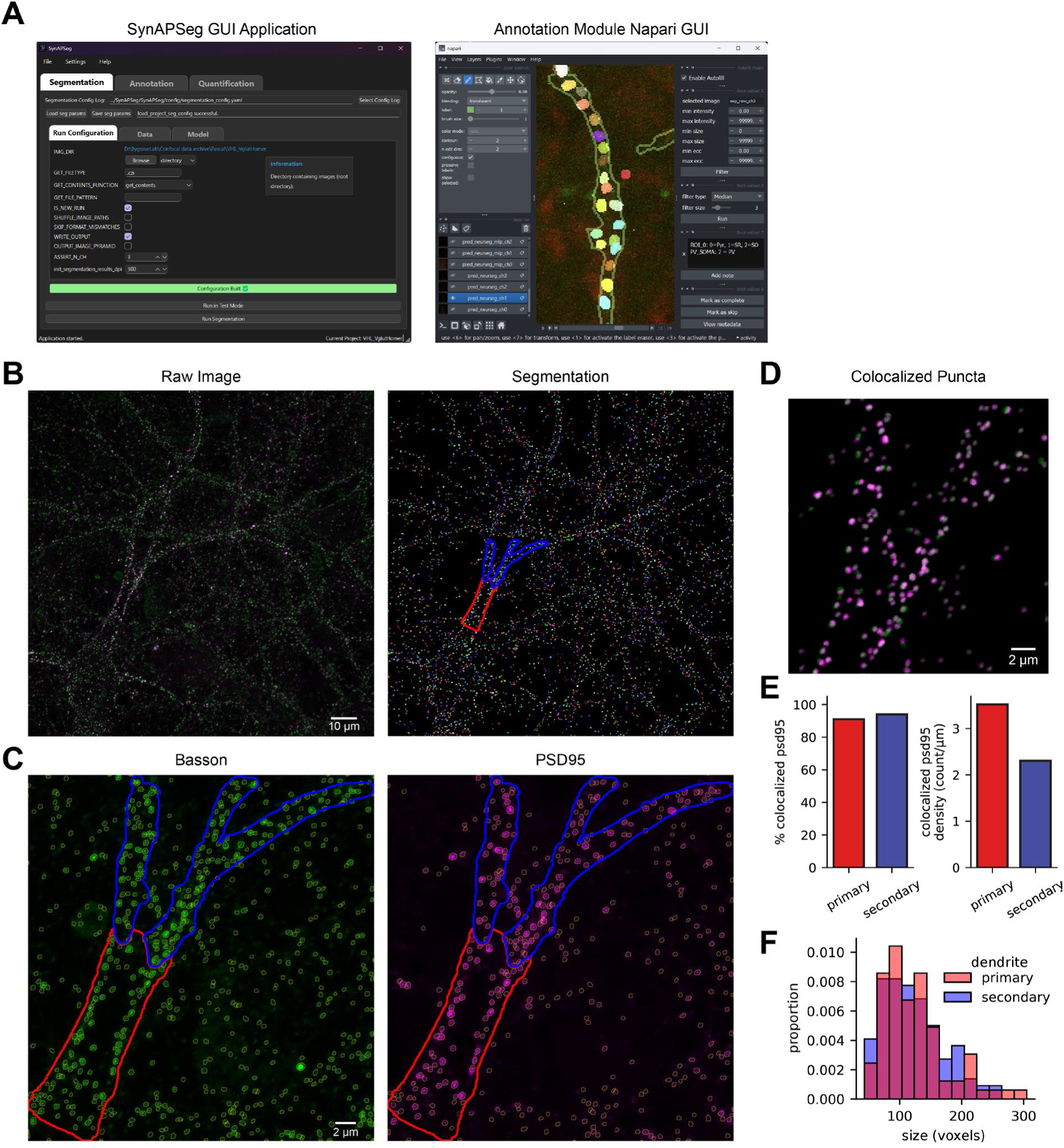
Implementation of the SynAPSeg framework for 3D colocalization analysis of pre-and postsynaptic markers. **A)** The SynAPSeg GUI provides an intuitive interface for configuring pipelines through interactive widgets. Setting up segmentation and quantification pipelines is accessible entirely through the GUI. Hovering over any parameter field provides detailed explanations. The Annotation Module utilizes an integrated Napari GUI enabling human-in-the-loop annotation and custom ROI definition facilitated by interactive image processing widgets (right). **B)** Maximum intensity z-projection of a Gad67+ cell (putative inhibitory neuron; Gad67 signal not shown) in neuronal culture where bassoon (green) and PSD95 (magenta) were labeled by immunocytochemistry (left). The resulting instance segmentation (right). **C**) High-magnification views of segmented bassoon (left) and PSD95 (right) puncta. Blue and red outlines indicate manually defined ROIs for primary and secondary dendritic segments, respectively. **D**) Visualization of colocalized puncta representing putative synapses identified via 3D object-based colocalization in SynAPSeg. **E**) Barplots (left) show the percentage of PSD95 puncta colocalized with bassoon (relative to the total number of PSD95 puncta) and the density of synapses (right) for primary and secondary dendrites. **F**) Feature extraction of individual synaptic objects, illustrated by the distribution of colocalized PSD95 puncta size (voxels) across dendritic ROIs.

**Supplemental Figure 5.**
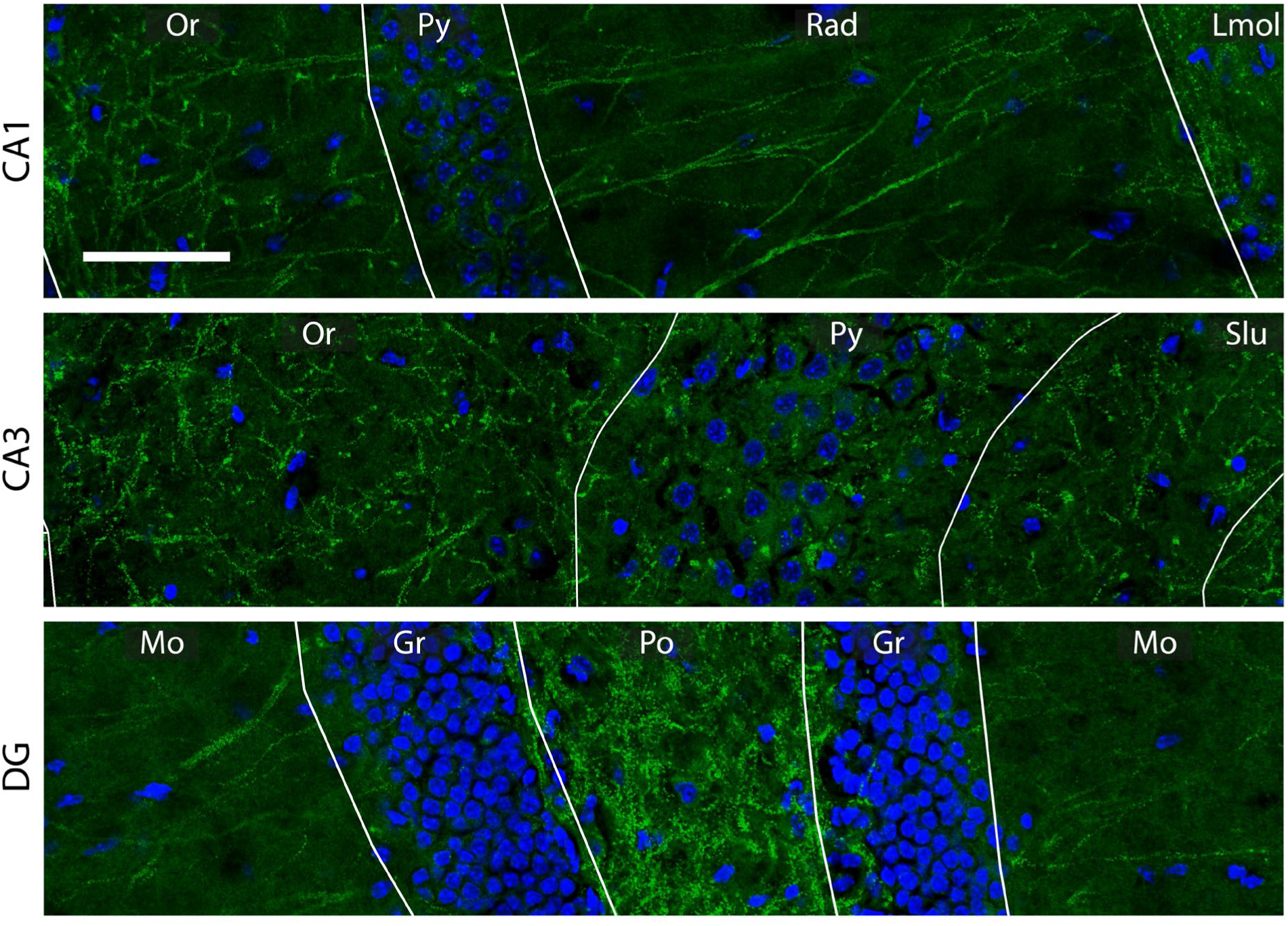
PSD95 puncta in the hippocampus. Representative images showing PSD95 puncta (green) and DAPI (blue) in subregions of CA1, CA3, and DG (top to bottom), illustrating qualitative differences in synaptic distribution and density. Scalebar: 50 µm. Hippocampal subregion abbreviations are defined as follows: Field CA1-3 (CA1-CA3), Oriens layer (Or), Pyramidal layer (Py), Radiatum layer (Rad), Lacunosum moleculare (Lmol), Stratum lucidum (Slu), Dentate gyrus (DG), Granule cell layer (Gr), Molecular layer (Mo), and Polymorphic layer (Po).

**Supplemental Figure 6.**
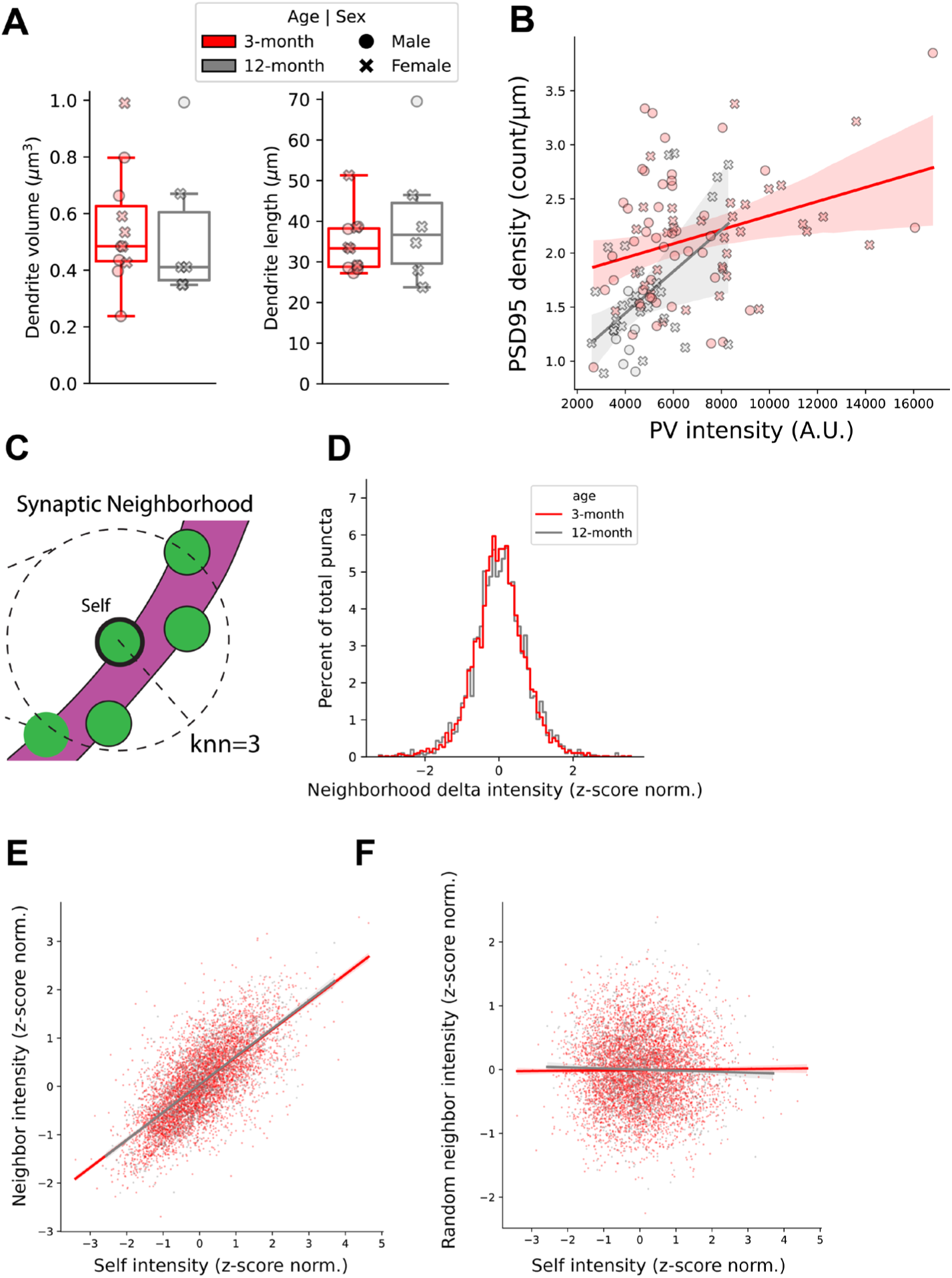
Extended morphological and spatial analysis of PV-IN dendrites and PSD95 puncta. **A)** Morphological properties of PV-IN dendrites analyzed. Boxplots show the total ROI area (µm^3^, left) and length (µm, right) for 3-month (red) and 12-month (gray) age groups. Data points represent animals. N = 11 3-month (N=5 females, 6 males) and 6 12-month: (5 females, 1 male). **B)** Correlation between PV-IN linear PSD95-GFP density and intensity of PV expression in dendritic segments. Individual points represent single dendritic segments; a positive correlation is observed in both age groups (Pearson; r=0.32, p < 0.01 3-month; r=0.51, p < 0.01 12-month). The correlations were not significantly different (Fisher r-to-z; z=1.07, p=0.29). **C-F)** Synaptic Neighborhood analysis. **C)** Schematic of the spatial nearest-neighbor analysis. The average z-scored intensity of the 3 closest synaptic neighbors was used to calculate “neighbor delta” (self-intensity minus average neighbor intensity) for each PSD95 puncta along individual dendrites. For all analyses raw intensity values of PSD95 puncta were z-score normalized within each dendrite independently **D)** Histogram showing the distribution of neighbor delta values; the y-axis displays the percentage of total puncta within each bin. **E)** Correlation between self-intensity and the average intensity of the surrounding synaptic neighborhood. A significant positive correlation is maintained across both age groups. (Pearson; r=0.70, p < 0.001 3-month; r=0.71, p < 0.001 12-month). The correlations were not significantly different (Fisher r-to-z; z=0.59, p=0.55) **F)** No correlation was observed between self-intensity and randomly shuffled neighbors (Pearson; r=0.01, p=0.49, 3-month; r=-0.03, p=0.24 12-month).

## Supplementary Tables

**Table S1.** Details of the SynAPSeg training dataset Table with the details related to sample preparation of imaging data used for training and validation of models.

|  | Dataset alias | nD | Marker | Labeling modality | Annotation type | Sample | Denoising | Z (um) | YX (um) |
| --- | --- | --- | --- | --- | --- | --- | --- | --- | --- |
| 0 | 2D_Tissue_vGatCre_PSD95GFP | 2D | PSD95-GFP | vgat-cre | ROI | tissue | n2v | 0.3 | 0.071 |
| 1 | 2D_Tissue_vGatCre_PSD95GFP | 2D | PSD95-GFP | vgat-cre | ROI | tissue | n2v | 0.3 | 0.071 |
| 2 | 2D_Tissue_vGatCre_PSD95GFP | 2D | PSD95-GFP | vgat-cre | ROI | tissue | none | 0.3 | 0.071 |
| 3 | 2D_Tissue_vGatCre_PSD95GFP | 2D | PSD95-GFP | vgat-cre | ROI | tissue | none | 0.3 | 0.071 |
| 4 | 2D_Tissue_vGatCre_PSD95GFP | 2D | PSD95-GFP | vgat-cre | ROI | tissue | none | 0.3 | 0.071 |
| 5 | 2D_Culture_Transfection_PSD95 | 2D | PSD95-FingR-GFP | transfection | full | ex vivo | none | 0.5 | 0.071 |
| 6 | 2D_Culture_Transfection_PSD95 | 2D | PSD95-FingR-GFP | transfection | full | ex vivo | none | 0.5 | 0.071 |
| 7 | 2D_Culture_Transfection_PSD95 | 2D | PSD95-FingR-GFP | transfection | full | ex vivo | none | 0.5 | 0.071 |
| 8 | 2D_Culture_Transfection_PSD95 | 2D | PSD95-FingR-GFP | transfection | full | ex vivo | none | 0.5 | 0.071 |
| 9 | 2D_Culture_Transfection_PSD95 | 2D | PSD95-FingR-GFP | transfection | full | ex vivo | none | 0.5 | 0.071 |
| 10 | 2D_Culture_Transfection_PSD95 | 2D | PSD95-FingR-GFP | transfection | full | ex vivo | none | 0.5 | 0.071 |
| 11 | 2D_Culture_Transfection_PSD95 | 2D | PSD95-FingR-GFP | transfection | full | ex vivo | n2v | 0.5 | 0.071 |
| 12 | 2D_Culture_Transfection_PSD95 | 2D | PSD95-FingR-GFP | transfection | full | ex vivo | none | 0.5 | 0.071 |
| 13 | 2D_Culture_Transfection_PSD95 | 2D | PSD95-FingR-GFP | transfection | full | ex vivo | none | 0.5 | 0.071 |
| 14 | 2D_Culture_Transfection_PSD95 | 2D | PSD95-FingR-GFP | transfection | full | ex vivo | none | 0.5 | 0.071 |
| 15 | 2D_Culture_Transfection_PSD95 | 2D | PSD95-FingR-GFP | transfection | full | ex vivo | none | 0.5 | 0.071 |
| 16 | 2D_Culture_Transfection_PSD95 | 2D | PSD95-FingR-GFP | transfection | full | ex vivo | none | 0.5 | 0.071 |
| 17 | 2D_Culture_IHC | 2D | Traf3 (1:400, Proteintech #66310-1-Ig) | IHC | ROI | ex vivo | none | 0.5 | 0.071 |
| 18 | 2D_Culture_IHC | 2D | Traf3 (1:400, Proteintech #66310-1-Ig) | IHC | ROI | ex vivo | none | 0.5 | 0.071 |
| 19 | 2D_Culture_IHC | 2D | Homer | IHC | ROI | ex vivo | none | 0.5 | 0.071 |
| 20 | 2D_Culture_IHC | 2D | Traf3 (1:400, Proteintech #66310-1-Ig) | IHC | ROI | ex vivo | none | 0.5 | 0.071 |
| 21 | 2D_Culture_IHC | 2D | Vglut1 (1:500, SYSY #135304) | IHC | ROI | ex vivo | none | 0.5 | 0.071 |
| 22 | 2D_Culture_IHC | 2D | Homer (SYSY #160002) | IHC | full | ex vivo | none | 0.5 | 0.071 |
| 23 | 2D_Culture_Transfection_PSD95 | 2D | PSD95-FingR-GFP | transfection | ROI | ex vivo | none | 0.5 | 0.085 |
| 24 | 2D_Culture_Transfection_PSD95 | 2D | PSD95-FingR-GFP | transfection | full | ex vivo | none | 0.5 | 0.085 |
| 25 | 2D_Culture_Transfection_PSD95 | 2D | Traf3-GFP | transfection | Full | ex vivo | none | 0.5 | 0.071 |
| 26 | 2D_Culture_Transfection_PSD95 | 2D | PSD95-mCh | transfection | full | ex vivo | none | 0.5 | 0.071 |
| 0 | 3D_Culture_transfection_PSD95 | 3D | PSD95-fingR | transfection | full | ex vivo | none | 0.19 | 0.071 |
| 1 | 3D_Culture_IHC | 3D | Traf3-GFP | IHC | ROI | ex vivo | none | 0.33 | 0.071 |
| 2 | 3D_Culture_IHC | 3D | Homer | IHC | ROI | ex vivo | none | 0.22 | 0.071 |
| 3 | 3D_Culture_IHC | 3D | Gephyrin (1:1000, SYSY #147011) | IHC | full | ex vivo | none | 0.4 | 0.071 |
| 4 | 3D_Culture_IHC | 3D | PSD95 | IHC | full | ex vivo | none | 0.4 | 0.071 |
| 5 | 3D_Tissue_vGatCre_PSD95GFP | 3D | PSD95-GFP | vgat-cre | full | ex vivo | none | 0.23 | 0.085 |
| 6 | 3D_Tissue_Geph | 3D | DIO-GephFingR-mScarlet | virus, vgat-cre | full | tissue | none | 0.28 | 0.1 |
| 7 | 3D_Tissue_vGatCre_PSD95GFP | 3D | PSD95-GFP | vgat-cre | full | tissue | n2v | 0.46 | 0.071 |
| 8 | 3D_Tissue_vGatCre_PSD95GFP | 3D | PSD95-GFP | vgat-cre | full | tissue | none | 0.46 | 0.071 |
| 9 | 3D_Tissue_vGatCre_P<br>SD95GFP | 3D | PSD95-GFP | vgat-cre | full | tissue | none | 0.46 | 0.071 |
| 10 | 3D_Tissue_vGatCre_P<br>SD95GFP | 3D | PSD95-GFP | vgat-cre | full | tissue | n2v | 0.46 | 0.071 |
| 11 | 3D_Tissue_vGatCre_P<br>SD95GFP | 3D | PSD95-GFP | vgat-cre | full | tissue | none | 0.46 | 0.071 |
| 12 | 3D_Tissue_EN_PSD95 | 3D | PSD95-GFP | aav-cre | ROI | tissue | none | 0.46 | 0.085 |
| 13 | 3D_Tissue_EN_PSD95 | 3D | PSD95-GFP | aav-cre,<br>PSD95-GFP<br>mouse | ROI | tissue | none | 0.46 | 0.085 |
| 0 | Benchmark1 | 2D | PSD95-FingR-GFP | transfecti<br>on | full | ex vivo | none | 0.5 | 0.071 |
| 1 | Benchmark2 | 2D | Vglut1<br>(1:500,<br>SYSY<br>#13530<br>4) | IHC | full | ex vivo | none | 0.5 | 0.071 |
| 2 | Benchmark3 | 3D | PSD95-GFP | vgat-cre | full | tissue | none | 0.46 | 0.071 |

**Table S2.** Details of statistical tests Table with statistical details for all tests performed and information about the N for each experiment.

| Main Figures | Test | Test statistics | P-value | Sample information |
| --- | --- | --- | --- | --- |
| 4E | **Two-Way ANOVA (Puncta density)** |  |  | N = 6 3-month (2 female 4 male) and 5 12-month (4 female 1 male) |
| | age * region | $F(11.0, 108) = 0.3175$ | 0.9807 | |
| | region | $F(11.0, 108) = 25.3367$ | < 0.0001 | |
| | age | $F(11.0, 108) = 0.4456$ | 0.5058 | |
|  | Šídák's multiple comparisons, paired T-test | df=5 |  |  |
|  | **Two-Way ANOVA (Puncta size)** |  |  |  |
| | age * region | $F(11.0, 108) = 0.1532$ | 0.9992 | |
| | region | $F(11.0, 108) = 14.5583$ | < 0.0001 | |
| | age | $F(11.0, 108) = 14.3116$ | 0.0003 | |
|  | Šídák's multiple comparisons, paired T-test | df=5 |  |  |
|  | **Two-Way ANOVA (Puncta intensity)** |  |  |  |
| | age * region | $F(11.0, 108) = 0.1608$ | 0.9990 | |
| | region | $F(11.0, 108) = 1.7421$ | 0.0736 | |
| | age | $F(11.0, 108) = 1.7438$ | 0.1894 | |
|  | Šídák's multiple comparisons, Welch's T-test | df=5 |  |  |
| 5C | Welch's T-test (PV intensity) | $t=-3.2447$, df=14.9854 | 0.0054 | N = 6 3-month (2 female 4 male) and 5 12-month (4 female 1 male) |
| 5E | Welch's T-test (PSD95 size) | $t=-0.6892$, df=6.2599 | 0.5154 | N = 6 3-month (2 female 4 male) and 5 12-month (4 female 1 male) |
| 5F | Mann Whitney test (PSD95 intensity) | $U=18$, df=6.7926 | 0.1490 | N = 6 3-month (2 female 4 male) and 5 12-month (4 female 1 male) |
| 5G | Welch's T-test (PSD95 linear density) | $t=-3.2562$, df=9.8126 | 0.0088 | N = 6 3-month (2 female 4 male) and 5 12-month (4 female 1 male) |
| Supplementary Figures |  |  |  |  |
| **S6A** | Mann Whitney test (Dendrite volume) | U=26, df=8.6464 | 0.5249 | N = 11 3-month (N=5 females, 6 males) and 6 12-month: (5 females, 1 male) |
| **S6A** | Mann Whitney test (Dendrite length) | U=38, df=6.0567 | 0.6605 | N = 11 3-month (N=5 females, 6 males) and 6 12-month: (5 females, 1 male) |
| **S6B** | Pearson correlation (PV intensity x PSD95 linear density) | 3-month r=0.3192 | 0.0039 | N = 80 3-month, and 33 12-month dendrites (dendrites per animal: 5-11, average: 7) |
|  |  | 12-month r=0.5084 | 0.0025 |  |
|  | Fisher r-to-z | z=1.0675 | 0.2857 |  |
| **S6E** | Pearson correlation (Nearest neighbor analysis of puncta intensity) | 3-month: r=0.7039 | <1E-300 | N = 5515 3-month and 1856 12-month PSD95 puncta |
|  |  | 12-month: r = 0.7119 | <1E-286 |  |
|  | Fisher r-to-z | z=-0.0700 | 0.9400 |  |
| **S6F** | Pearson correlation (Randomly sampled neighbors) | 3-month: r=0.0094 | 0.4859 | N = 5515 3-month and 1856 12-month PSD95 puncta |
|  |  | 12-month: r=-0.0275 | 0.2359 |  |

## Notes

https://doi.org/10.5281/zenodo.18988899

### Competing Interest Statement

The authors have declared no competing interest.

